# The mutational features of aristolochic acid-induced mouse and human liver cancers

**DOI:** 10.1101/507301

**Authors:** Zhao-Ning Lu, Qing Luo, Li-Nan Zhao, Yi Shi, Xian-Bin Su, Ze-Guang Han

## Abstract

Aristolochic acid (AA) derived from traditional Chinese herbal remedies has recently been statistically associated with human liver cancer; however, the causal relationships between AA and liver cancer and the underlying evolutionary process of AA-mediated mutagenesis during tumorigenesis are obscure. Here, we subjected mice, including *Pten*-deficient ones, to aristolochic acid I (AAI) alone or a combination of AAI and carbon tetrachloride (CCl_4_), which may induce liver injury. Significantly, AAI promoted the development of liver cancer, including hepatocellular carcinoma and intrahepatic cholangiocarcinoma, in a dose-dependent manner, and it increased the incidence of liver cancer, together with CCl_4_ or *Pten* deficiency. AAI could lead to DNA damage and AAI-DNA adducts that initiate liver cancer via characteristic A>T transversions, as indicated by the comprehensive genomic analysis, which revealed recurrent mutations in *Hras* and some genes encoding components of the Ras/Raf, PI3K, Notch, Hippo, Wnt, DNA polymerase family and the SWI/SNF complex, some of which are also often found in human liver cancer. Mutational signature analysis across human cancer types revealed that the AA-related dominant signature was especially implicated in liver cancer in China, based on very stringent criteria derived from the animal cancer form, in which mutations of *TP53* and *JAK1* are prone to be significantly enriched. Interestingly, AAI-mediated characteristic A>T mutations were the earliest genetic event driving malignant subclonal evolution in mouse and human liver cancer. In general, this study provides documented evidence for AA-induced liver cancer with featured mutational processes during malignant clonal evolution, laying a solid foundation for the prevention and diagnosis of AA-associated human cancers, especially liver cancer.

## Introduction

Aristolochic acid (AA) is present in plants in the genera *Aristolochia, Bragantia, Asarum* and others^1^, which have been widely used in traditional Chinese herbal remedies. AA is one of the most potent carcinogens known to man, belonging to the Group I human carcinogens classified the by International Agency for Research on Cancer (IARC). Aristolochic acid I (AAI) and ? (AAII) are the major components of the AA mixture contained in the plant extract of *Aristolochia* species^2^. AA is a genotoxic carcinogen because its metabolite can bind purines to form AA-DNA adducts, aristolactam (AL)-DNA adducts (dA-AL and dG-AL), which are specific markers of exposure to aristolochic acids and induce DNA mutations with characteristic adenine-to-thymine (A>T) transversions *in vitro* and *in vivo*^2,3^.

The dA-AL-I (7-(deoxyadenosin-N6-yl) aristolactam I) adducts induced by AAI can show long-term persistence in renal tissue^4^, which may have led to the occurrence of aristolochic acid nephropathy (AAN) in Belgian women who had taken weight-reducing pills containing *Aristolochia fangchi*^5^ and Balkan endemic nephropathy (BEN) through dietary contamination with *Aristolochia clematitis* seeds^6^. Both nephropathies are associated with urothelial carcinoma because AA-DNA adducts have been found in kidney tissue and urothelial tumor tissues of patients with AAN or BEN^3,5,6^. In Taiwan, approximately one-third of the people consume Chinese herbal remedies containing AA, which could be associated with the highest incidence of upper urinary tract cancers (UTUC) in the world^7^. The genome-wide mutational signature of characteristic A>T transversions (COSMIC signature 22), specifically reflecting AA-implicated mutagenesis, is frequently found in Taiwanese UTUC^7,8^.

Recently, AA has been statistically associated with human liver cancer. We found, for the first time, that the characteristic A:T to T:A transversions were significantly enriched in 4 of 10 (40%) hepatitis B virus (HBV)-associated hepatocellular carcinoma (HCC) specimens from China, indicative of AA exposure in HCC tumorigenesis^9^. A survey in larger cohorts of HCC patients indicated that the AA-implicated mutational signature was discovered in some Asian HCC patients, especially in more than 75% of Taiwanese HCC cases^10^.

There has been no direct evidence that AA can induce liver cancer until now, although AA-DNA adducts have been detected in many organs, including the liver, in experimental animals exposed to AA during a relatively short peroid^11^, and even in the livers of some nephropathy patients with known AA exposure^12,13^. To confirm whether AA can directly induce liver cancer, including HCC, here we subjected mice, including *Pten*-deficient ones, to AAI alone or a combination of AAI and carbon tetrachloride (CCl_4_), a well-documented liver injury agent. Significantly, AAI administration alone increased the incidence of liver cancer in a dose-dependent manner, and the combination of AAI and CCl_4_ also led to a higher incidence of mouse liver cancer. Interestingly, the types of liver cancer included HCC, intrahepatic cholangiocarcinoma (ICC), and combined hepatocellular and intrahepatic cholangiocarcinoma (cHCC-ICC). Genome-wide analysis of the AAI-induced liver cancer showed the characteristic mutational signature and process during clonal evolution, providing new insights into the pathogenesis of AA-induced liver cancer.

## Results

### AAI can induce mouse liver cancer

To validate whether AA could directly induce liver cancer, especially HCC, we first subjected C57BL/6 male mice to AAI administration alone by intraperitoneal injection. Based on previous research^11,14^, AAI was administered at a lower dose (2.5 and 5 mg/kg body weight) for injection of 3, 7, and 14 times, respectively (see **Methods**). Moreover, considering the possibility that AAI could enhance tumorigenesis due to liver injury or a genetic defect, we designed the combination of AAI and CCl_4_ to treat mice, in which CCl_4_ can induce liver injury, compensatory proliferation, inflammation, and fibrosis^15^; we also subjected liver-specific *Pten*-deficient mice, who frequently develop liver cancer by 74–78 weeks of age^16^, to AAI administration. In general, a total of eight experimental groups of mice subjected to AAI administration (**Fig. 1a** and **Supplementary Fig. 1a**) included the following: (I) “AAI (3x)”, administration of AAI at a dose of 2.5 mg/kg every other day for 3 doses at 2 weeks after birth; (II) “AAI (14x)”, administration of AAI at a dose of 2.5 mg/kg/day for 14 days at 2 weeks of age; (III) “AAI (high 3x)”, administration of AAI at a dose of 5 mg/kg every other day for 3 doses at 2 weeks of age; (IV) “AAI (high 14x)”, administration of AAI at a dose of 5 mg/kg/day for 14 days at 2 weeks of age; (v) “AAI (3x) + CCl_4_”, administration of CCl_4_ three times per week for 4 weeks at 2 months after AAI injection; (VI) “AAI (14x) + CCl_4_”, administration of CCl_4_ once per week for 10 weeks at 4 weeks after AAI injection; (VII) “AAI (7x)”, administration of AAI at a dose of 2.5 mg/kg every other day for 7 doses at 1 week of age, to observe the effect of AAI on younger fetal livers; (VIII) “AAI (high 14x, *Pten*^*LKO*^)”, administration of AAI at a dose of 5 mg/kg/day for 14 days at 2 weeks of age in liver-specific *Pten*-deficient mice. In addition, male mice were injected with CCl_4_ alone, which was administered once or three times per week at a dose of 0.5 ml/kg body weight, and vehicle as the control group (**Supplementary Fig. 1a**).

**Figure 1.**
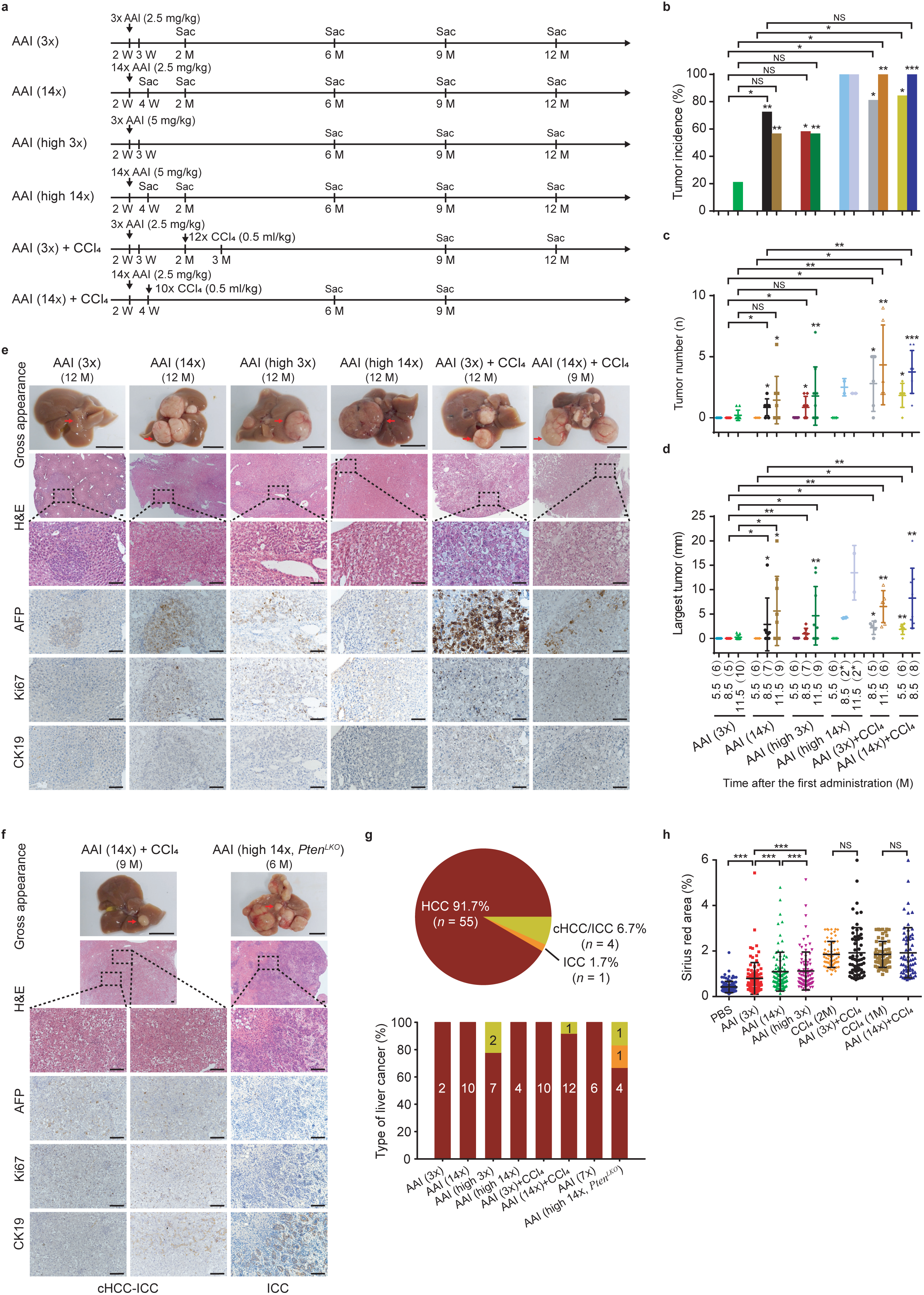
AAI can induce liver cancer. (**a**) Simplified diagram of liver cancer induction in C57BL/6 male mice with AAI alone or a combination of AAI and CCl_4_, where the dosages and time points of drug administration are indicated by arrows, and samples are harvested at the indicated time of sacrifice mice (Sac). (**b-d**) Tumor incidence (**b**), tumor number (**c**), and largest tumor size (**d**) of AAI-induced liver cancer. The numbers in parentheses are the numbers of mice in the corresponding group. The numbers of mice in the control groups corresponding to the first four experimental groups in the figure were 6 (5.5 M), 10 (8.5 M), and 12 (11.5 M); that in the fifth group were 5 (CCl_4_ (2 M), 8.5 M) and 6 (CCl_4_ (2 M), 11.5 M); and that in the sixth model were 6 (CCl_4_ (1 M), 5.5 M) and 6 (CCl_4_ (1 M), 8.5 M). The asterisk directly above each group indicates a significant difference compared with the corresponding control group. The numbers marked with an asterisk indicate the numbers of surviving mice, and the initial number of mice in each group was 11. (**e, f**) Representative images of gross appearance (scale bars, 1 cm), hematoxylin and eosin (H&E) staining (scale bars, 100 μm), and immunohistochemistry (IHC) analysis with anti-AFP, Ki67 and CK19 antibodies (scale bars, 100 μm) of AAI-induced HCCs (**e**), cHCC-ICC and ICC (**f**). (**g**) Proportion of types of HCC, cHCC-ICC and ICC in all C57BL/6 male mice with liver cancers (up) and each group (down). The numbers in the column charts indicate the number of mice with the corresponding cancer types in each group. (**h**) Quantification of PicroSirius Red histochemistry staining in liver slices from the different experimental groups and control groups. The numbers of mice corresponding to each group were 12 (time after first administration, 11.5 M), 10 (11.5 M), 9 (11.5 M), 9 (11.5 M), 5 (8.5 M), 5 (8.5 M), 6 (5.5 M) and 6 (5.5 M). (**b-d, h**) Values indicated by long and short horizontal lines represent the mean ± SD. Asterisks signify significant differences using the two-sided Student’s *t*-test or Wilcoxon rank-sum test and Fisher’s exact test. \**P* < 0.05; \*\**P* < 0.01; \*\*\**P* < 0.001; NS, not significant.

Interestingly, liver cancer occurred in all eight experimental groups of AAI administration (**Fig. 1b-g** and **Supplementary Fig. 1b-p**). AAI administration alone significantly promoted the development of liver cancer in a dose-dependent manner. A lower dosage of AAI (“AAI (3x)”) led to liver cancer development in 2 (20%) out of 10 mice at 11.5 months after the first AAI administration, demonstrating a statistically increased incidence compared with the control group without AAI treatment (*P* = 0.038) (**Supplementary Fig. 1b**). Significantly, a greater number of injections (“AAI (14x)” *vs.* “AAI (3x)”) was associated with earlier occurrence (8.5 M *vs.* 11.5 M) and larger tumor sizes (11.5 M, mean: 5.62 mm *vs.* 0.22 mm, *P* = 0.048) of liver cancer (**Fig. 1d**). Under the same AAI administration durations, the larger the dosage (“AAI (high 3x)” *vs.* “AAI (3x)”), the larger was the tumor size (11.5 M, mean: 4.63 mm *vs*. 0.22 mm, *P* = 0.048) (**Fig. 1d**). However, many mice in the “AAI (high 14x)” group died during the experimental observation period (**Supplementary Fig. 1q**), and all 4 surviving mice at 8.5 or 11.5 months after the first AAI administration developed liver cancer (**Supplementary Fig. 1j**). In addition, the mice in the “AAI (7x)” injection group at the age of 1 week also displayed an increment in tumor incidence, number and size, compared to the “AAI (14x)” group, although this difference was not statistically significant (**Supplementary Fig. 1b-d, o**).

Compared with AAI administration alone, the combination of both AAI and CCl_4_ led to a significantly higher incidence of mouse liver cancer. The combined models (“AAI (3x) + CCl_4_ “*vs.* “AAI (3x)”; “AAI (14x) + CCl_4_ “ *vs.* “AAI (14x)”) resulted in an earlier tumor occurrence (8.5 M *vs.* 11.5 M; 5.5 M *vs.* 8.5 M), higher tumor incidence (11.5 M, 100% *vs*. 20%, *P* = 0.007; 8.5 M, 100% *vs.* 71.4%, *P* = 0.2), greater number of tumor nodules (11.5 M, mean: 4.3 *vs.* 0.2, *P* = 0.00048; 8.5 M, mean: 3.75 *vs.* 0.86, *P* = 0.0013) and larger tumor sizes (11.5 M, mean: 6.53 mm *vs*. 0.22 mm, *P* = 0.00051; 8.5 M, mean: 8.25 *vs*. 2.86, *P* = 0.014) (**Fig. 1b-d**).

Moreover, compared with the same genetic background mice as a control, the liver-specific *Pten*-deficient mice (*Pten*^*LKO*^) treated with AAI alone developed liver cancer (6 M, 100% *vs.* 0%, *P* = 0.002) (**Supplementary Fig. 1b-d**), along with obvious bile duct hyperplasia in adjacent liver tissues (**Supplementary Fig. 1p**).

Among the examined 84 livers from the above mice that received AAI administration, 60 mice (71.4%) developed liver cancer. We checked these tumors based on the microscopic morphology and immunohistochemistry staining for Ki67, a proliferative index; α-fetoprotein (AFP), a well-known HCC marker; and cytokeratin 19 (CK19), a cholangiocyte marker. Interestingly, 55 (91.7%) of 60 tumors were observed to be HCCs, which exhibited expansive growth, hyperchromatic and enlarged nuclei, an increased nuclear-to-cytoplasmic ratio, a high Ki67 proliferative index, an absence of normal liver architecture, and focal expression of AFP (**Fig. 1e, g** and **Supplementary Fig. 1b-p**). Additionally, 4 (6.7%) of 60 tumors were classified as combined HCC and intrahepatic cholangiocarcinoma (cHCC-ICC) because both AFP and CK19-positive cells were present in the same tumors (**Fig. 1f, g**), which were obtained from different groups treated with AAI alone, a combination of AAI and CCl_4_, and liver-specific *Pten*-deficient mice, respectively (**Fig. 1g and Supplementary Fig. 1h, i, n, p**). Interestingly, 1 (1.7%) *Pten*^*LKO*^ mouse developed ICC with CK19-positive cells in an examined tumor nodule (**Fig. 1f, g**); however, the HCC nodule was also observed in the same liver (**Supplementary Fig. 1p**). It should be pointed out that no visible liver tumors were detected in any of the control groups.

In addition to liver cancer, AAI also promoted liver fibrosis in a dose-dependent manner. The minimum amount of AAI administration (“AAI (3x)” group) also led to fibrillar collagen deposition, as detected by Sirius red staining in noncancer livers, compared to the controls without AAI treatment (Sirius red area: 0.79% *vs.* 0.42%, *P* = 9.9 × 10 ^-10^) (**Fig. 1g** and **Supplementary Fig. 1r**). The mice in the “AAI (14x)” and “AAI (high 3x)” groups displayed a profound increment of fibrosis compared with those in the “AAI (3x)” group (Sirius red area: 1.81% *vs*. 0.79%, *P* = 3.2 × 10 ^-15^; 1.12% *vs*. 0.79%, *P* = 1.44 × 10 ^-4^, respectively) (**Fig. 1h** and **Supplementary Fig. 1r**). Interestingly, the degree of liver fibrosis paralleled the incidence of liver cancer with AAI administration alone.

Moreover, we also checked the other organs of these mice treated with AAI. Hydronephrosis or renal cysts were found in the mice that were administered AAI (**Supplementary Fig. 1s**). However, no visible tumors were found in other organs, such as lung, spleen, stomach, ureter, bladder and testis.

The collective data indicated that AAI could result in liver cancer, including HCC, ICC and cHCC-ICC, in a dose-dependent fashion; when the liver was injured or displayed a genetic defect such as *Pten* deficiency, AAI could synergistically promote liver cancer tumorigenesis. The above data also implied that AAI could trigger genetic lesions in liver progenitor cells with bipotent potential towards hepatocytes or cholangiocytes, which further develop three subtypes of liver cancer under the genetic differentiation program.

### AAI causes DNA damage and dA-AL-I adducts in liver

It is known that AA is a genotoxic agent that can form DNA adducts such as dA-AL and dG-AL; however, whether AA causes genomic DNA damage in liver cells is unclear. We first examined the phosphorylated histone γ-H2AX, a biomarker for DNA double-strand breaks, in the livers of mice with AAI administration alone (“AAI (14x)”), through immunofluorescence staining. Interestingly, the phosphorylated γ-H2AX level was obviously increased in the liver at 1 month of age (four days after completion of AAI administration) (**Fig. 2a**); however, during the subsequent 2-12 months, the γ-H2AX level was reduced in liver (**Supplementary Fig. 2a**). Excluding the phosphorylated γ-H2AX level, we further evaluated the p53 level and its downstream target molecule Bax as a cellular response to DNA damage, in these livers via Western blotting, which revealed that both γ-H2AX and p53 levels were markedly increased in these livers (**Fig. 2b**), along with a slight upregulation of Bax. These data suggested that AAI could give rise to DNA damage.

**Figure 2.**
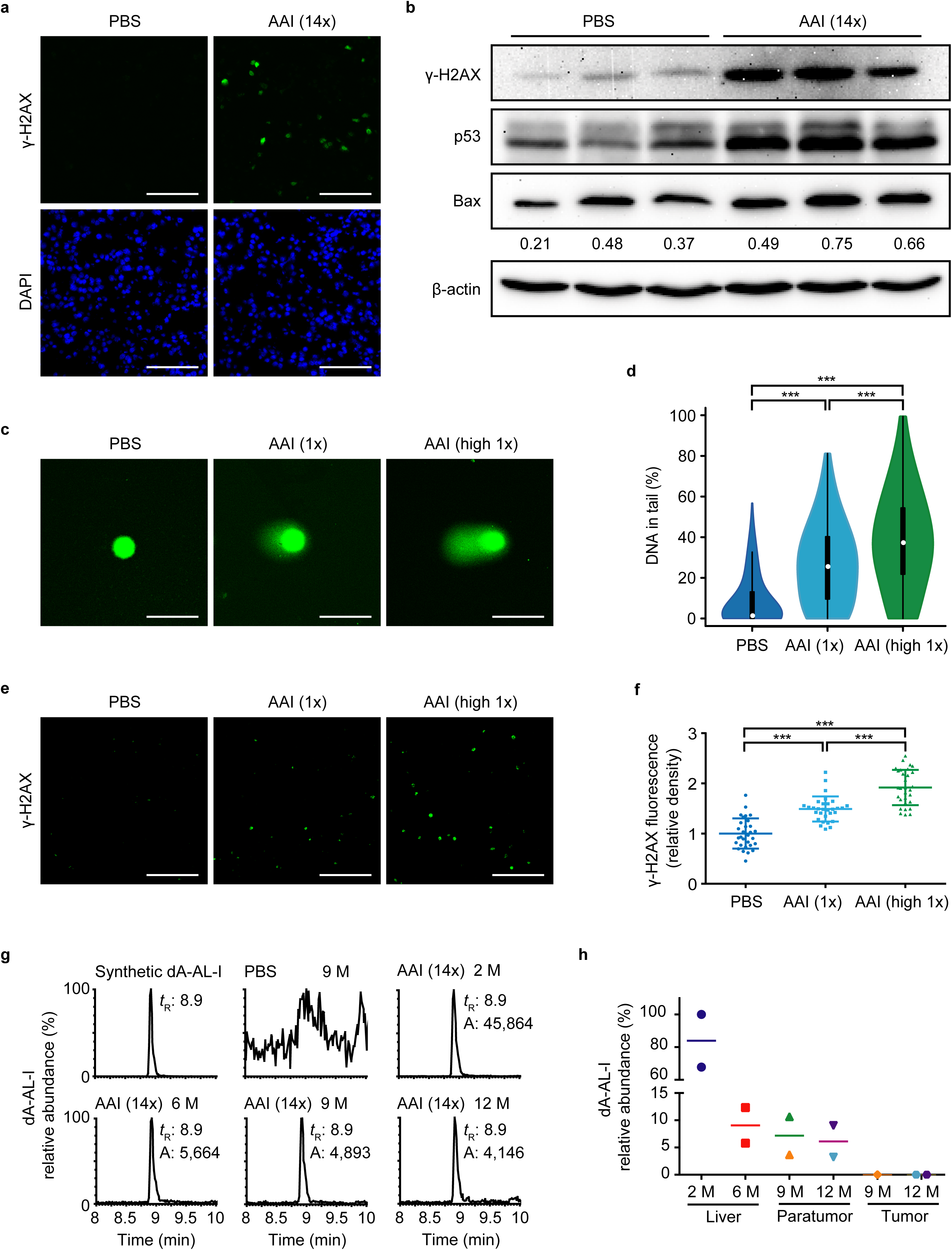
AAI can cause liver DNA damage. (**a**) The γ-H2AX level was measured by immunofluorescence assay in livers from the control group (PBS) and “AAI (14x)” mice at 1 month of age. (**b**) γ-H2AX, p53 and Bax levels were measured by Western blotting assay (n = 3) in mouse livers from the control group (PBS) and “AAI (14x)” group. The numbers under the Bax band are relative intensity values. (**c, d**) The DNA strand breaks were measured by the alkaline comet assay in liver cells from 2-week-old mice at 3 h once after PBS or AAI (2.5 mg/kg or 5 mg/kg) injection, including representative images (**c**) and quantitative analyses (**d**, the number of mice per group is 4; the number of nuclei per group: 600, at least 100 nuclei from each mouse). The white dot, thick black bar in the center and thin black line extended from it in (**d**) stand for the median, interquartile range and 95% confidence intervals. (**e, f**) The γ-H2AX level was measured by the immunofluorescence assay in livers from the above samples, including representative images (**e**) and quantitative analyses (**f**, the number of mice per group: 3; 10 nonoverlapping fields at ×400 magnification per mouse). Values indicated by long and short horizontal lines represent the mean ± SD. (**g, h**) The relative abundance (*m/z* 427) of dA-AL-I was measured by MS in livers of “AAI (14x)” mice at the indicated ages, including representative images (**g**) and quantitative analyses (**h**). Spots with the same colors indicate the paratumors and tumors are from the same mouse. The horizontal lines in (**h**) denote the mean. Asterisks signify significant differences using the two-sided Student’s *t*-test or Wilcoxon rank-sum test. \*\*\**P* < 0.001. Scale bars, 100 μm.

To confirm whether AAI could indeed lead to DNA damage in liver, we employed the alkaline comet assay to directly detect DNA strand breaks in livers 3 hours after AAI administration with 2.5 mg and 5 mg/kg dosages, respectively. The data demonstrated that AAI could cause DNA strand breaks in mouse livers in a dose-dependent fashion (**Fig. 2c, d**), in parallel with the increased phosphorylated γ-H2AX level via immunofluorescence staining (**Fig. 2e, f**), which was positively correlated with the incidence of liver cancer. Except for the increased γ-H2AX level in an exposure time-depend manner (**Supplementary Fig. 2b, c**), phosphorylated ATR, a molecule that responds to DNA damage, was upregulated in liver at 12 h after AAI administration (**Supplementary Fig. 2d**). These data revealed that AAI directly triggered DNA damage in mouse livers.

AA is known to form AA-DNA adducts that are further processed to form somatic mutations through infidelity DNA repair system, which could be critical step in tumorigenesis. We thus examined the AAI-mediated dA-AL-I adduct in mouse livers after AAI administration by mass spectrometry, using the identified synthetic dA-AL-I as a reference (**Supplementary Fig. 2e, f**). Significantly, we could detect dA-AL-I adducts in all examined noncancerous livers from “AAI (14x)” mice (**Fig. 2g**), and the quantity of the adduct in these livers gradually decreased along the different time points after AAI administration (**Fig. 2g, h**), while the quantity of dA-AL-I in kidneys was generally higher than in livers from the same mice (**Supplementary Fig. 2g**). However, we could not detect the dA-AL-I adducts in three matched liver cancers (**Fig. 2h**), implying that, within these tumor cells, the activated DNA repair system had removed the adduct or the adduct could be diluted by repeated DNA replication via cell cycle progression.

The above data indicated that AAI caused DNA damage, including DNA double-strand breaks, and dA-AL-I adducts in liver cells, which triggered the cellular response and DNA repair system. This process could further lead to genomic instability and somatic mutations that contribute to tumorigenesis.

### AAI leads to the characteristic mutational signature of A to T transversions

To survey the genomic instability and somatic mutations triggered by AAI, we performed whole-genome sequencing (WGS) for DNA copy number variations (CNVs), whole-exome sequencing (WES) and transcriptome analysis of 11 AAI-induced liver tumor nodules, three matched adjacent noncancerous livers, three livers prior to the occurrence of tumors (from the “AAI (3x)” group) and two livers from mice treated with CCl_4_ alone, in which their corresponding mouse tails for sequencing were used as the reference controls (**Supplementary Table 1**). Among the 11 tumor nodules, 3 were respectively resected from 3 mice of the “AAI (14x)” group (labeled the AAI group), and the other 8 tumors from another 3 mice in the “AAI (3x) + CCl_4_” group (labeled as combination group) (**Supplementary Table 1** and **Supplementary Fig. 1g, l**), where 3 and 4 discrete tumor nodules were resected respectively from two mice (**Supplementary Fig. 3a**).

CNV analysis of the 11 tumor nodules by WGS at the depth of about 3-fold, compared to their corresponding tail tissues as references, showed that the AA-induced tumors barely had obvious CNV alterations (**Supplementary Table 2**). WES at the average depth of 267-fold for all examined 11 tumor nodules and 8 nontumor livers from the AAI, CCl_4_ and combination groups, in which WES data of 62-fold for their corresponding tail tissues were references, identified a total of 8107 single-nucleotide variants (SNVs) and 704 small insertions and deletions (indels) in the 11 tumor nodules (**Supplementary Tables 1 and 3**).

The somatic SNVs and indels of tumors from the AAI group were significantly more abundant than those in the combination group (mean: 1555 *vs.* 518, *P* = 0.012) (**Supplementary Table 1**), possibly because of the larger AAI dosage in “AAI (14x)” group than in the combination group. Interestingly, somatic mutations were also found in the paratumor livers and noncancerous livers with or without tumors (mean: 130 *vs.* 76, *P* = 0.036) (**Supplementary Table 1**) of mice treated with AAI or CCl_4_ alone, respectively, suggesting that the increased somatic mutations in livers could be prerequisite to liver cancer development triggered by AAI.

Significantly, the tumors exhibited remarkably high proportions (69%) of A>T transversions, whereas the nontumor liver tissues, except for one (M4P: 33%), did not show such feature (10%) (**Supplementary Fig. 3b, c** and **Supplementary Table 4**). Notably, the load of A>T mutations in the AAI group was larger than those in the combination group (mean: 1043 *vs*. 305, *P* = 0.012) (**Supplementary Table 4**), which was consistent with the observation that the total applied AAI amount was higher in the AAI group. The mutational profile of each tumor nodule was depicted (**Fig. 3a** and **Supplementary Fig. 3d-g**) and showed that the trinucleotide context of the highest proportion of A>T mutations was CTG (or CAG on the complementary strand). The pentanucleotide context of the highest proportion of A>T mutations was CCTGT (or ACAGG on the complementary strand) (**Supplementary Fig. 3h**). In the series of mutational deciphering analysis, 7 COSMIC mutational signatures were detected (**Fig. 3b**), of which signature 22 related to AA was obviously dominant in all tumor nodules. Except for signature 22, signatures 1, 5, 6, 17 and 23 were detected in these mouse tumor nodules, of which signature 1 related to deamination of 5-methylcytosine, and 5 related to aging, have been found in all human cancer types and most cancer samples, while signature 6 is associated with defective DNA mismatch repair, and the etiologies of signatures 5, 17 and 23 remains unknown. Tobacco-associated signature 4 was surprisingly detected in two mouse tumors.

**Figure 3.**
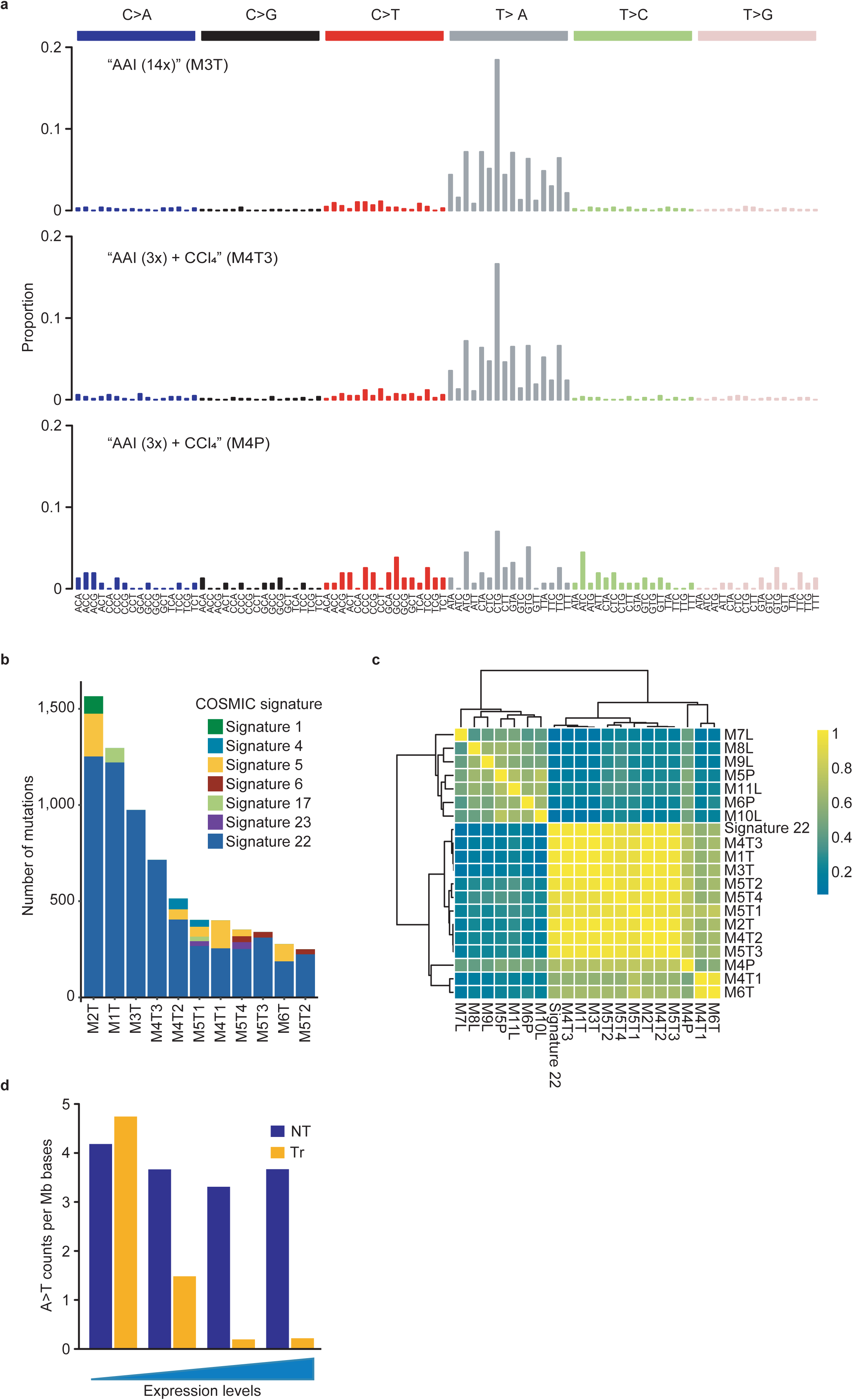
Mutational signatures of AAI-induced mouse liver cancer. (**a**) Representative trinucleotide contextualized mutational spectra in AAI-induced liver cancer. (**b**) Estimated COSMIC mutational signature contributions for each mouse liver cancer. Mutational signature decomposing with known liver cancer signatures (i.e., COSMIC signatures 1, 4, 5, 6, 12, 16, 17, 22, 23 and 24) was performed using the least square root algorithm. The AA signature (or COSMIC signature 22) was dominant throughout the mouse liver cancer. (**c**) Cosine similarities of trinucleotide mutational spectra between the tumor/noncancerous liver samples and the AA signature. M1T, M2T, M3T, M4T1, M4T2, M4T3, M5T1, M5T2, M5T3, M5T4 and M6T refer to AAI-induced liver cancer; M4P, M5P and M6P are the paratumor liver tissues of the mice in the combination group; M7L, M8L and M9L are the livers from the “AAI (3x)” group. M10L and M11L are the liver tissues from the CCl_4_ -treated group. (**d**) The mutational frequency of A>T transversions in transcribed and nontranscribed regions per megabase (Mb) in these genes as a function of the expression level. The genes with expression were divided into 4 expression quintiles according to the expression levels. NT, nontranscribed strand; Tr, transcribed strand.

However, it was noteworthy that two tumors, M4T1 and M6T, presented higher levels of T > G mutations (marked with a prominent peak at ATG > AGG) (**Supplementary Fig. 3e**), the etiology of which remains unknown. Subsequently, the cosine similarities between the mutational spectra of these tumors and signature 22 (typical AA signature) were calculated. Except for M4T1 and M6T, which showed a somewhat lower similarity (0.59 and 0.64) to signature 22 due to distortion of the higher T > G mutations, the mutational profiles of the tumors were nearly identical to signature 22, having a cosine similarity larger than 0.9 (**Fig. 3c** and **Supplementary Table 5**).

Interestingly, the mutational spectrum of one paratumor tissue (M4P, A>T, 33%) also exhibited a similar feature to signature 22 (cosine similarity = 0.69) (**Fig. 3a**), which was obviously higher than the other nontumor liver tissues (average cosine similarity = 0.14) (**Fig. 3c** and **Supplementary Table 5**). This result indicated that M4P could be associated with a precancerous process, which was consistent with the pathology of hyperplasia (**Supplementary Fig. 3a**), along with higher somatic mutations (189) than the other two paratumor livers from the same group (mean: 100.5), albeit being significantly lower than those of the tumor nodules from the same group (mean: 518) (**Supplementary Table 1**).

Except for M4P, other nontumor liver tissues exhibited C>T (average 36%) rather than A>T (average 10%) as the predominant mutation category (**Supplementary Fig. 3c** and **Supplementary Table 4**). Their mutational spectra exhibited higher similarities to signatures 5 and 6 (**Supplementary Fig. 3i** and **Supplementary Table 5**).

Previous studies have revealed that AA-induced mutations are likely to be transcriptionally strand biased^8^. Here, the calculated average ratio of A>T mutations on the nontranscribed strand versus the transcribed-strand was 2.02 (*P* < 0.001) in tumors (**Supplementary Fig. 3j** and **Supplementary Table 6**), indicating the existence of a transcription-coupled repairing (TCR) mechanism to fix the AA-mediated mutations. To further validate the influence of the transcription history on the asymmetries of the A>T strand distribution, we investigated the A>T mutation counts on both strands in the five defined gene expression categories, from low to high expressions, according to the gene expression profiles of the 11 tumor nodules (**Supplementary Table 7**). Next, the mutation counts on the transcribed versus nontranscribed strands were analyzed within each defined gene category, showing that the strand bias of A>T mutations was indeed positively correlated with the gene expression levels (**Fig. 3d**).

The collective data revealed that the characteristic mutational signature of A to T transversions and the COSMIC signature 22 induced by AAI were involved in tumorigenesis and could be necessary and critical for the development of liver cancer.

### Affected driver genes and signaling pathways

The AAI-mediated characteristic A>T mutations could damage the driver genes that could initiate liver cancer. To identify the driver mutations of these genes in AAI-induced mouse tumors, we searched for genes that were mutated more frequently than expected given the average observed mutation frequency. Interestingly, we found 1919 genes with nonsynonymous mutation in the 11 tumor nodules (**Supplementary Tables 1**), of which 98 genes with a total of 123 nonsynonymous somatic mutations belong to the Cancer Gene Census as known driver genes (Tier 1 for 77 genes), or those with strong indications for a role in cancer but with less extensive available evidence (Tier 2 for 21 genes) in human cancers (**Supplementary Table 8**). Interestingly, 92 (75%) of the 123 nonsynonymous mutations were A>T mutations.

The statistically significantly mutated genes included *Hras, Sfi1, Muc4, Sp140, Vmn2r121* and *Inpp5d* (**Supplementary Table 9**). The well-studied oncogenic A>T mutations led to the change of *Hras* Q61L (CAA>CTA) in 8 of 11 (72.7%) tumor nodules, and *Kras* Q61L (CAA>CTA) and *Braf* V637E (GTG>GAG) were also identified in two other tumor nodules (**Fig. 4a**), indicating that the cancer-promoting mutations of the Ras/Raf pathway were crucial in AAI-induced liver cancer. Interestingly, 4 of 11 (36.4%) tumors presented *Muc4* (4 A>T mutations), in which the same *Muc4* (c.3869T>A) mutation was detected in two tumors (**Fig. 4a**). *Muc4* as an oncogene is listed in the Cancer Gene Census Tier 2, mutations of have appeared in many human cancers, including HCC^10,17,18^, and are associated with tumor metastasis^19^. *Sfi1* encoding a spindle assembly associated protein, which was reported to be mutated in human HCC^18^, showed 8 mutations (A>T mutations) in 4 mouse tumors (**Fig. 4a**). *Sp140* encodes a member of the SP100 family of proteins, *Inpp5d* encodes a member of the inositol polyphosphate-5-phosphatase (INPP5) family, and unknown functional *Vmn2r121* also showed a higher mutation frequency in these mouse tumors (3/11), of which *Inpp5d* (3/11), also named *SHIP1*, involved in the PI3K-AKT pathway, has been known to be mutated in human cancers, including HCC^17^ (**Supplementary Table 3** and **Fig. 4a**).

**Figure 4.**
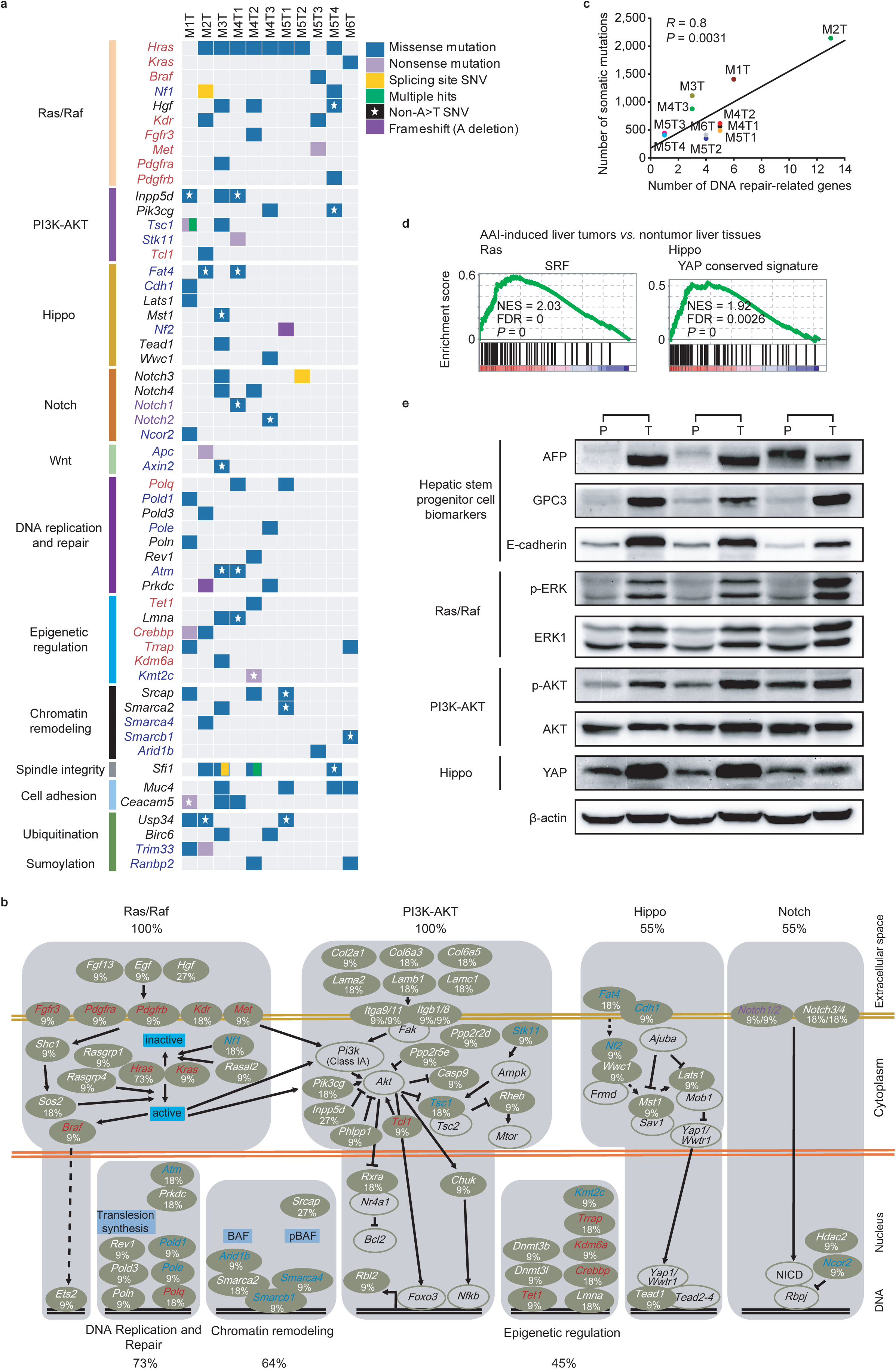
The genes and signaling pathways affected by AAI-mediated mutations. (**a**) The categories of the statistically and empirically important genes with somatic mutations in liver cancer. The genes in red or blue refer to the proto-oncogenes and tumor-suppressor genes listed in COSMIC Cancer Gene Census Tier 1, respectively. Genes in purple font refer to driver genes without a clear definition in terms of proto-oncogenes and tumor-suppressor genes. (**b**) Major signaling pathways involving genetic alterations in AAI-induced mouse liver cancer. A brown background indicates mutated genes; a white background denotes unmutated genes. Genes in red and blue refer to proto-oncogenes and tumor-suppressor genes listed in COSMIC Cancer Gene Census Tier 1, respectively. Those in purple refer to driver genes without a clear definition in terms of proto-oncogenes and tumor-suppressor genes. Percentages stand for the proportion of gene or genes in the pathway altered in liver cancer. (**c**) Correlations between the number of DNA repair-related genes with the total mutation counts in mouse liver tumors. (**d**) Gene set enrichment analysis (GSEA) plot of SRF and YAP motif target gene sets. SRF is a key regulatory transcription factor in the Ras signaling pathway, and YAP is a key regulatory transcription factor in the Hippo signaling pathway. (**e**) AFP, GPC3, E-cadherin, p-ERK, ERK1, p-AKT, AKT and YAP levels were measured by Western blotting assay (n = 3) in mouse paratumors (“P”) and tumors (“T”) from “AAI (3x) + CCl_4_” (18 M) group.

Both the Ras/Raf and PI3K-AKT pathways could participate in the pathogenesis of all these liver tumors (**Fig. 4a, b**). Except for *Ras* and *Braf*, five genes that regulate Ras activity, *Nf1* (2/11), *Rasal2* (1/11), *Sos2* (1/11), *Rasgrp1* (1/11) and *Rasgrp4* (1/11), were also identified with A>T mutations (**Supplementary Table 3**). In addition, some genes encoding growth factors and receptors with tyrosine kinase, such as *Hgf* (3/11), *Egf* (1/11), *Fgf13* (1/11), *Kdr* (2/11), *Pdgfra* (1/11), *Pdgfrb* (1/11), and *Fgfr3* (1/11), except for *Met* (1/11), were also influenced by A>T mutations. *Hgf* mutations also appear in human HCC^10,17,18^. Moreover, the mutant genes were significantly enriched in the PI3K-AKT signaling pathway (*P* = 6.6 × 10 ^-6^) (**Supplementary Table 10** and **Fig. 4a, b**), which included those encoding growth factors and the receptors mentioned earlier. Except for *Inpp5d*, some genes encoding phosphoinositide-3-kinase (PI3K), such as *Pik3cg* (2/11), and modulators of AKT activity such as *Ppp2r2d* (1/11), *Ppp2r5e* (1/11), *Phlpp1* (1/11) and *Tcl1* (1/11), were also influenced by A>T mutations. The tumor suppressor gene *Tsc1* (2/11), as a negative regulator of mTORC1 and *Rheb* (1/11) activating the protein kinase activity of mTORC1, demonstrated A>T mutations. (**Fig. 4a**).

Some mutations could damage development-related genes, including components of the Hippo, Notch and Wnt pathways (**Fig. 4a, b**). *Fat4* (2/11), *Cdh1* (1/11), *Nf2* (1/11), *Lats1* (1/11), *Mst1* (1/11), *Tead1* (1/11) and *Wwc1* (1/11), belonging to the Hippo signaling pathway, had A>T mutations. *Notch1* (1/11), *Notch2* (1/11), *Notch3* (2/11), *Notch4* (2/11) and *Ncor2* (1/11), encoding components of the NOTCH signaling pathway, had A>T mutations. Three genes involved in the WNT signaling pathway, *Apc* (1/11), *Axin2* (1/11) and *Wnt1* (1/11), were mutated in 2 tumors (**Supplementary Table 3**).

It was noticeable that these genes encoding DNA polymerases, including *Polq* (2/11), *Pold1* (1/11), *Pold3* (1/11), *Pole* (1/11), *Poln* (1/11) and *Rev1* (1/11), had A>T mutations in 6 of 11 (54.5%) tumors (**Fig. 4a, b**). Excluding DNA replication, these DNA polymerases perform exonucleolytic proofreading for DNA repair. It is known that defective DNA polymerase proofreading contributes to human malignancy, and DNA polymerase mutations in the exonuclease domain have been reported in human tumors with an extremely high mutation load^20,21^. Other DNA repair-related genes, such as *Atm* (2/11), *Prkdc* (2/11), *Mcm8* (2/11) and *Trp53bp1* (1/11), were also mutated in these tumors (**Supplementary Table 3**). Here, we statistically analyzed the correlation between somatic mutations and these gene mutations, showing that the mutations of these DNA genes in tumors were positively associated with the total somatic mutations (**Fig. 4c**).

Some genes related to epigenetic regulation exhibited somatic mutations (**Fig. 4a, b**), including *Tet1* (1/11), *Dnmt3b* (1/11) *and Dnmt3l* (1/11) for DNA methylation, *Crebbp* (2/11), *Trrap* (2/11), *Kdm6a* (1/11) and *Kmt2c* (1/11) for histone modifications and *Srcap* (3/11), *Smarca2* (2/11), *Smarca4* (1/11), *Smarcb1* (1/11) and *Arid1b* (1/11) for the chromatin remodeling SWI/SNF complex. Mutations of these genes have been described in human cancers, including liver cancer.

In addition, some genes related to ubiquitination and sumoylation were also mutated, including *Usp34* (3/11), *Trim33* (2/11), *Birc6* (2/11) and *Ranbp2* (2/11) (**Fig. 4a**). *Usp34* encoding ubiquitin carboxyl-terminal hydrolase 34 can remove conjugated ubiquitin from Axin1 and Axin2, as a regulator of the Wnt signaling pathway, which is also mutated in human HCC^10,17,18^.

To further assess the effect of these mutations on the pathogenesis of liver cancer, we analyzed the transcriptome data from these tumor nodules. Some target genes of important pathways were upregulated, such as the Ras, PI3K-AKT, Hippo and Wnt pathways disrupted by the mutations, especially Ras and Hippo (**Supplementary Fig. 3k, Fig. 4d and Supplementary Table 11**). Interestingly, some genes, such as *Afp, Dlk1, Gpc-3, Prom1, Itga6, Cd34* and *Igdcc4* related to stem cells/progenitor cells, along with downstream target genes, such as *Fstl1, Dab2, Hes1* and *Mycn* of the Hippo, Notch and Wnt pathways, were upregulated in liver tumors, suggesting that cell differentiation arrest or dedifferentiation occurred in these liver cancers (**Supplementary Fig. 3k** and **Supplementary Table 11**). The transcription of cell cycle-related genes, such as *Ccnd1, Ccne1*, and *Cdks,* were increased in tumors, possibly due to activation of the Ras and PI3K-AKT pathways (**Supplementary Table 7**). Further, the activation of these signaling pathways including Ras, PI3K-AKT and Hippo was verified in these AAI-induced tumors, as compared to adjacent non-tumorous livers (**Fig. 4e** and **Supplementary Fig. 3l**), along with the up-regulated hepatic stem cells/progenitor cell biomarkers.

The collective data suggested that AAI contributed to tumorigenesis of liver cancer through activating the RAS pathway, in combination with other deregulated important pathways such as PI3K-AKT, DNA replication and repair, the chromatin remodeling SWI/SNF complex, epigenetic regulation, the development-related Hippo, Notch and Wnt pathways, spindle integrity, and cell adhesion.

### AAI-mediated mutations are the early event during malignant clonal evolution

Though it was testified that the AA signature was dominant in mouse liver tumors, we had particular interest in whether the AA-mediated mutations were the originating source driving tumor initiation and progression, especially for tumors in the combination group composed of AAI and CCl_4_, as the application of CCl_4_ inevitably cast doubts on the role of AA in the carcinogenic processes. Therefore, we further investigated the clonal architecture and AA-related mutational signature distribution in these malignant subclones within tumors from the “AAI alone” and “AAI and CCl_4_ combination” groups.

We first performed a clonality analysis of these 11 liver tumor nodules (see **Methods**). Ten of the 11 nodules exhibited multiple subclones, of which 3 contained 3 subclones and 7 had 2 subclones (**Fig. 5a, b** and **Supplementary Fig. 4a-f, left)**.

**Figure 5.**
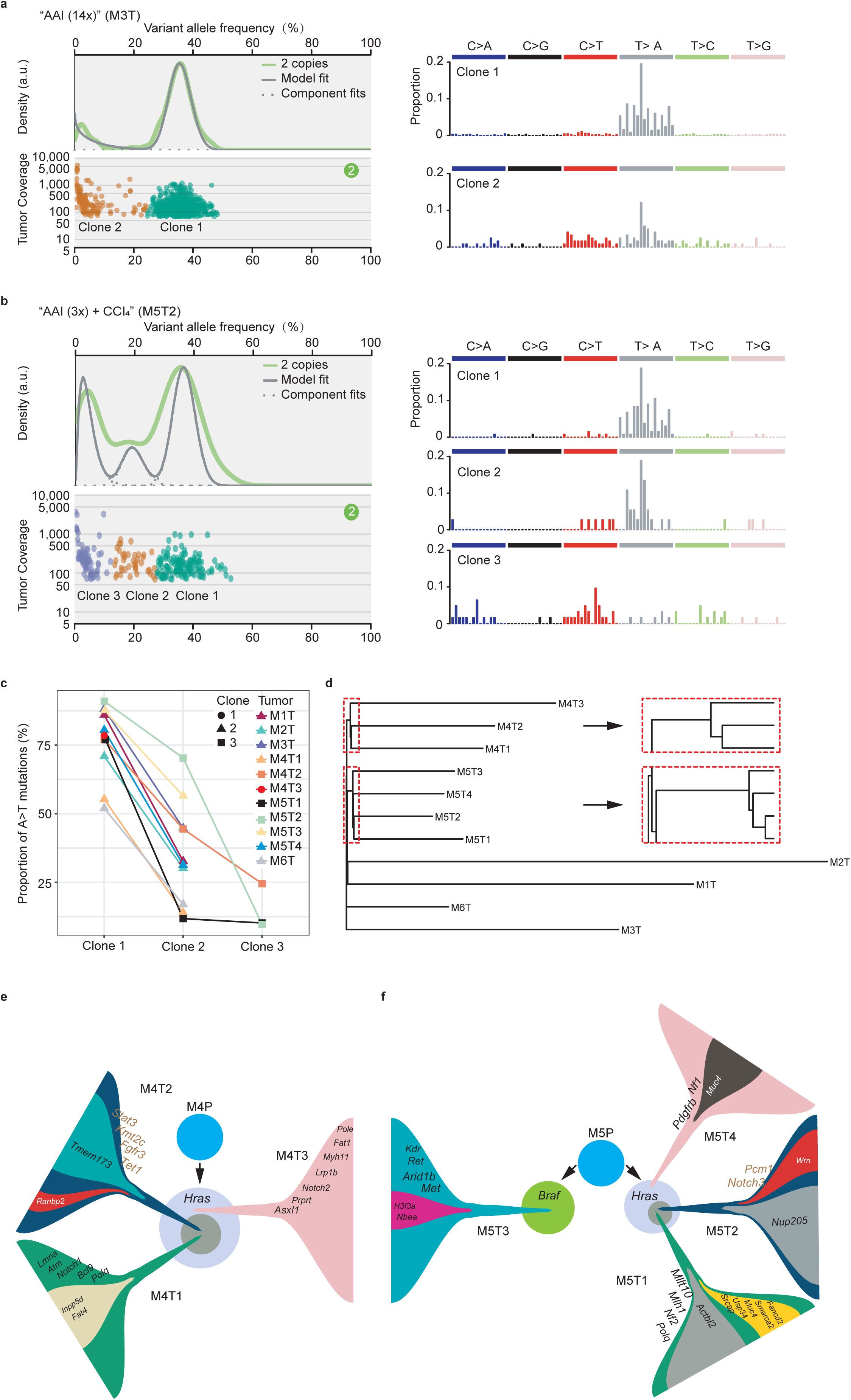
Clonal architecture and phylogenetic reconstructions of AAI-induced mouse liver cancer. (**a, b, left**) Malignant clonal architecture reconstructions within M3T1 and M5T2 tumors. Each peak indicates one subclone. The subclones lying at the right end had the largest mutational allele frequency and therefore represent the founding clones. Others are subclones. (**a, b, right**) Trinucleotide mutational spectra of the founding clone and subclones within M3T1 and M5T2. The A>T transversions were predominant in the earliest founding clones and diminished in the later formed subclones. (**c**) The downward-pointing lines of A>T mutations in different subclones within these 11 tumor nodules. The A>T mutation proportions deposited in each clone in the multiclonal tumors. (**d**) The phylogenetic tree was reconstructed using the neighbor-joining algorithm with the R package ape (v5.2). (**e**) A common ancestry evolutionary model for three discrete tumor nodules within M4 liver. The font size of the genes reflects the allele frequency in each tumor. (**f**) Common ancestry evolutionary model for three discrete tumor nodules within M5 liver. The font size reflects the genes allele frequency in each tumor.

Pure tumor was expected to present a high-density region with nearly a 50% variant allele frequency (VAF) in the Sciclone deconvolution results^22^. The estimated weights of VAF in the tumor dominant clones, however, ranged from 16% to 39% (**Supplementary Table 12**), reflecting a substantial immunological cell infiltration into the tumors as presented in the pathological sections (**Supplementary Fig. 3a**). Alternatively, the subclones could have been initiated in parallel style in the tumorigenesis procedures, especially in tumors M1T, M4T1 and M6T, as their dominant clone had VAF centered at 16%, 19% and 17%. If the subclones formed simultaneously, they should present similar mutational signatures induced by the same etiologies. However, we found that the mutational signatures varied within the multiple subclones within each tumor (**Fig. 5a, b, right** and **Supplementary Fig. 4a-f, right**). In addition, there was a trend towards a diminished AA signature from the higher-weighted to the lower-weighted clones in their signature profiles. Therefore, we considered the subclones with the largest weight to be the initiating founding subclones and the lower-weighted subclones to be formed sequentially in later processes ^22^. To determine how the AA-related mutations evolved between the founding clone and subsequent subclones, we calculated the A>T proportions of each subclone within the tumors. For example, after characterizing the multiclonal architectures in M3T (two subclones) and M5T2 (three subclones) (**Fig. 5a, b, left**), we then retrieved their mutational profiles (**Fig. 5a, b, right**). Significantly, the results indicated that the AAI-mediated A>T transversions were predominant in the earliest founding clones and then gradually were reduced in the later subclones. The other 8 tumor nodules composed of multiple subclones also exhibited a similar AA-related signature distribution pattern **(Supplementary Fig. 4a-f).** This result indicated that the AAI-mediated A>T mutations were the early event during the malignant clonal evolution process, contributing to an average of 82% in the AAI group (M1T, 86%; M2T, 71%; M3T, 88%) and an average of 75% in the combination group (M4T1, 55%; M4T2, 77%; M5T1, 78%; M5T2, 91%; M5T3, 88%; M5T4, 80% and M6T, 52%). By contrast, the non-A>T mutational patterns, such as C>T mutations, slightly increased gradually during clonal evolution and merged into the late subclones **(Fig. 5a, b and Supplementary Fig. 4 a-f, right)**. As illustrated by the downward-directed lines in these 11 tumor nodules (**Fig. 5c**), there was a general trend of A>T mutations that diminished along with the malignant clonal progression within tumors, typically M5T1 and M5T2, in which the A>T transversions almost disappeared in their late-formed subclones.

Based on the above analysis, we speculate that, regardless of the AAI alone group or the combination group, AAI-mediated A>T somatic mutations are responsible for the initiation of liver cancer, and the second non-A>T mutations drive malignant clonal evolution and tumor progression, possibly through a synergistic effect between AAI-mediated A>T and non-A>T mutations.

### AAI-mediated tumors exhibit diversiform evolution process

The above clonality analysis depicted the intratumor clonal heterogeneity within single tumor nodules, but AAI can induce multiple tumor nodules in the same livers in some mice, and the phylogenetic relationship of these nodules is unclear. To explore their phylogenetic relationship and evolutionary process, here we investigated these tumor nodules and their paratumor livers from mice in the combination group, with a total of the 8 discrete tumor nodules; 3 were from one mouse (M4), 4 were from another (M5), and 1 was from the last mouse (M6). We reconstructed their phylogenetic tree to examine the relationship of the somatic mutational patterns among the discrete tumor nodules within the same mice. To establish a control, we applied the reconstruction to all 11 tumor nodules to generate their phylogenetic tree (**see Method**). It was seen that, despite a tiny overlapping distance between M1T and M2T, the other tumor nodules were categorized properly, in accordance with their mouse source (**Fig. 5d**), implying that multiple nodules in M4 and M5 could have evolved from identical ancestors, respectively.

Interestingly, the three nodules in M4 shared the specific oncogenic *Hras* Q61L mutation and another 15 identical mutations, supporting the assumption that the separate tumor nodules might have originated from a common ancestor. Moreover, 12 passenger mutations were shared by both the paratumor liver and all three tumor nodules, although they had lower allelic frequencies in the paratumor tissue (e.g., Chr14: 5140816 C>G), suggesting that the tumor could be initiated by the emerging driver mutations, such as *Hras* Q61L, in the background originating cell with passenger mutations (**Fig. 5e**). Additionally, M4T1 and M4T2 might branch later than M4T3, as indicated in the phylogenetic tree (**Fig. 5d**).

Unlike the tumor nodules in mouse M4, the four tumor nodules in M5 did not share a commonly known driver mutation, although three (M5T1, M5T2 and M5T4) of them shared an identical oncogenic *Hras* Q61L mutation, and the other (M5T3) harbored an oncogenic *Braf* V637E mutation. However, as indicated in the phylogenetic tree, all four nodules could have arisen from the same ancestor and then evolved separately in the late phase. The observation that M5T1, M5T2 and M5T4 were located within a branch (**Fig. 5d**), rather than M5T3, revealed a closer phylogenetic relationship, in accordance with their differences in the initiating driving force. Interestingly, all four tumor nodules in M5 also shared 10 somatic mutations with their paratumor liver, including Chr11: 3176625 G>A that increased the mutation allele frequency from the paratumor liver to the tumor, suggesting that these tumor nodules could be initiated from the same precancerous cells through the emerging driver mutations such as *Hras* Q61L or *Braf* V637E (**Fig. 5f**). Although all four tumor nodules could originate from the same precancerous cells with a similar genetic background, the M5T3 nodule with the *Braf* V637E mutation was distinguished from the other three nodules sharing a common ancestor, wherein the two malignant transformed clones had undergone parallel evolution within M5 liver (**Fig. 5f**).

To reveal the kinship between different subclones within the separated tumor nodules from M4 and M5, here we adopted the assumption that the second subclone was generated dependently from the founding clones. However, whether the third weighted subclone was generated dependently or independently of the second-weighted clone was unclear (**Supplementary Fig. 4g**). Here, the M4T2, M5T1 and M5T2 nodules were composed of three subclones (**Supplementary Fig. 4d, e**), where the third subclones within single nodules were depicted to emerge via parallel evolution along with the second subclones by driver genes such as *Ranbp2, Actbl2, Smarca2* and others (**Fig. 5e, f**). The relationship between the second and third clones could also be replaced by the other model as provided in Supplementary Fig. 4g with the same suggested driver genes and cell proportions.

In M6, both the paratumor liver and the corresponding tumor had 30 overlapping somatic mutations, among which 6 expanded their allelic frequencies more than 5 times from the paratumor liver to the tumor. Next, we examined the presence of mutated reads of essential genes in the paratumor liver tissue in Integrated Genome Viewer (IGV). We found that the paratumor liver of M6 has the same positioned *Muc4* mutation reads in its corresponding tumor, albeit with a very low number (2 reads) (**Supplementary Fig. 4h**). Therefore, we speculate that the *Muc4* mutation, along with the other passenger mutations (like the 6 expanded mutations), was not sufficient to trigger tumor initiation and that one of the *Muc4* mutated cells, if acquiring the oncogenic *Kras* mutation, would be malignantly transformed and then become proliferative (**Supplementary Fig. 4i**).

Together with the above clonality analysis within tumor nodules (**Fig. 5e, f** and **Supplementary Fig. 4i**), we may see that, except for Ras/Raf A>T mutations as the earliest events in the tumorigenesis of M4, M5 and M6 mice, the patterns of other driver mutations exhibit obvious heterogeneity among different nodules and subclones within single nodules. The different driver genes, such as *Polq, Fgfr3, Met, Asxl1, Pdgfrb, Notch3, Mllt10* and *Tet1* with A>T mutations and late emerging driver genes such as *Fat4, Smarca2* and *Inpp5d* with non-A>T mutations, synergistically facilitate malignant subclonal evaluation.

### Human liver cancer exhibits an AA-mediated mutational signature

To explore the AA signature intensities in human cancers, we first grasped a quick estimation of the AA signature contribution through the webserver mSignatureDB (**see Methods**). Interestingly, some human cancer resources presented a possible characteristic A>T mutational signature to certain degrees (**Supplementary Fig. 5a**). However, current approaches for signature deconvolution were mostly based on nonnegative matrix factorization (NMF), which do not consider mutation counts. Based on a simulated dataset with 844 samples (see **Methods**), we noticed that, when the mutation number fell within 100, the mean squared error (MSE) of deconvolution increased exponentially as the mutation number decreased (**Supplementary Fig. 5b**). Therefore, in an effort to reduce false positives with a low number of mutation counts, we improved the traditional decomposing strategy by introducing the bootstrap sampling technique to make up for the shortcoming that the deconvolution originally did not provide empirical *P* values. Afterwards, we used the simulated dataset to evaluate the performance of the bootstrap performance by considering its accuracy, specificity, sensitivity and F1 measure. Surprisingly, the evaluation revealed no false positive mistakes (specificity = 1) throughout each threshold (**Supplementary Fig. 5c**). In addition, the method yielded fine-tuned accuracies, sensitivities and F1 measures for detecting the AA signature intensity in the interval between 10 to 90%. Therefore, we decided to identify the AA signature intensities in human cancers with a cutoff of both 0 and 10% (*P* = 0.05). Furthermore, we required that the AA signature proportion should be larger than the MSE to balance the instabilities due to low mutation counts and unpredictable noise. Using this method, we found that the primary positive detection of AA signature in the majority of tumors and leukemia was prone to be false.

However, with this rigorous method, we identified an AA signature in liver cancer, including HCC and ICC (**Table 1**). We detected a characteristic AA signature in 52 (20%) of 313 China (mainland) HCC samples catalogued in the International Cancer Genome Consortium (ICGC) project, 68 (69%) of 98 Taiwan-based samples^10^, and 7 (8%) of 88 samples that were accepted in Hong Kong^23^, as well as 6 (55%) of another 11 samples in mainland China^24^. In total, 133 (26%) of 510 Chinese HCCs were identified with an AA signature. Moreover, we detected the AA signature in 3 (< 1%) of 594 HCCs from Japan^17,18^, 22 (10%) of 231 HCCs in Korea^25^, 4 (44%) of 9 HCCs in Singapore^26^, 29 (10%) of 364 HCCs in the US from The Cancer Genome Atlas (TCGA) dataset, and 1 (< 1%) of 249 HCCs from France in the ICGC. Among the TCGA HCCs, Asian ethnicity patients had higher detection rate of AA signature, which is 24 (15%) of 160. In addition, we noticed that 11 (11%) of 103 ICC from China^27^ showed the AA signature, indicating that AA might play a role in the etiology of human ICCs (**Table 1**).

**Table 1.** **Aristolochic acid exposure in human liver cancers**

| Cancer type | Data source | Regions | Number of patients | Numbers (%) of patients with AA exposure inferred by different criteria |  |  |  |
| --- | --- | --- | --- | --- | --- | --- | --- |
|  |  |  |  | > 0* | > 10** | Nonsilent mutations in known driver genes | > 52%*** |
| HCC | ICGC | China (mainland) | 313 | 52 (17) | 51 (17) | 27 (9) | 12 (4) |
|  | Kan, *et al.*<sup>23</sup> | China (Hongkong) | 88 | 7 (8) | 4 (5) | 1 (1) | 1 (1) |
|  | Lin, *et al.*<sup>24</sup> | China (mainland) | 11 | 6 (55) | 6 (55) | 2 (18) | 2 (18) |
|  | Ng, *et al.*<sup>10</sup> | Taiwan | 98 | 68 (69) | 68 (69) | 53 (54) | 42 (43) |
|  | Chinese HCC total |  | 510 | 133 (26) | 129 (25) | 83 (16) | 57 (11) |
|  | TCGA | USA (Asian) | 160 | 24 (15) | 23 (14) | 13 (8) | 4 (3) |
|  | TCGA | USA (others) | 204 | 5 (2) | 4 (2) | 3 (1) | 0 (0) |
|  | ICGC | France | 249 | 1 (<1) | 0 (0) | 0 (0) | 0 (0) |
|  | ICGC<sup>17,18</sup> | Japan | 594 | 3 (<1) | 1(<1) | 2 (<1) | 0 (0) |
|  | Zhai, *et al.*<sup>26</sup> | Singapore | 9 | 4 (44) | 4 (44) | 1 (11) | 0 (0) |
|  | Ahn, *et al.*<sup>25</sup> | Korea | 231 | 22 (10) | 22 (10) | 11 (5) | 3 (1) |
|  | Worldwide HCC total |  | 1957 | 192 (10) | 183 (9) | 113 (6) | 64 (3) |
| ICC | Zou, *et al.*<sup>27</sup> | China (mainland) | 103 | 11 (11) | 11 (11) | 4 (4) | 3 (3) |

Note: \* indicates the estimated lower boundary of 95% confidence interval of AA signature exposure larger than 0 ( $P < 0.05$ ); \*\* indicates the estimated lower boundary of 95% confidence interval of AA signature exposure larger than 10% ( $P < 0.05$ ); \*\*\* indicates the estimated lower boundary of 95% confidence interval of AA signature exposure larger than 52% ( $P < 0.05$ ).

We also investigated other human cancer types. Bladder cancer in China^28^ and kidney cancer in Europe (from ICGC) presented different extents of AA signature contributions (**Supplementary Table 13)**. In addition, it was noteworthy that we identified 1 case of esophagus cancer in China (from ICGC) that exhibited the AA signature (**Supplementary Tables 13, 14)**. Generally, in all the cancer types, liver cancer, including HCC and ICC, presents the most disturbingly high proportions of the AA signature. Additionally, we retrieved the T>A mutations of liver cancer in COSMIC and found that the T>A mutation profile was highly consistent with the AA signature (cosine similarity = 0.94) (**Supplementary Fig. 5d**), which implicated that the AA signature operated predominantly in causing the T>A mutation in human liver cancers. Next, we compared the AA signature intensities in the affected human cancers, which exhibited wide intertype and regional variabilities (**Fig. 6a**). Kidney cancer and liver cancer were more susceptible to AA genotoxicity, as their AA intensities were prominently higher. According to the proportions of affected populations and their AA signature intensities, it seemed that the most influenced cancer type was HCC in China.

**Figure 6.**
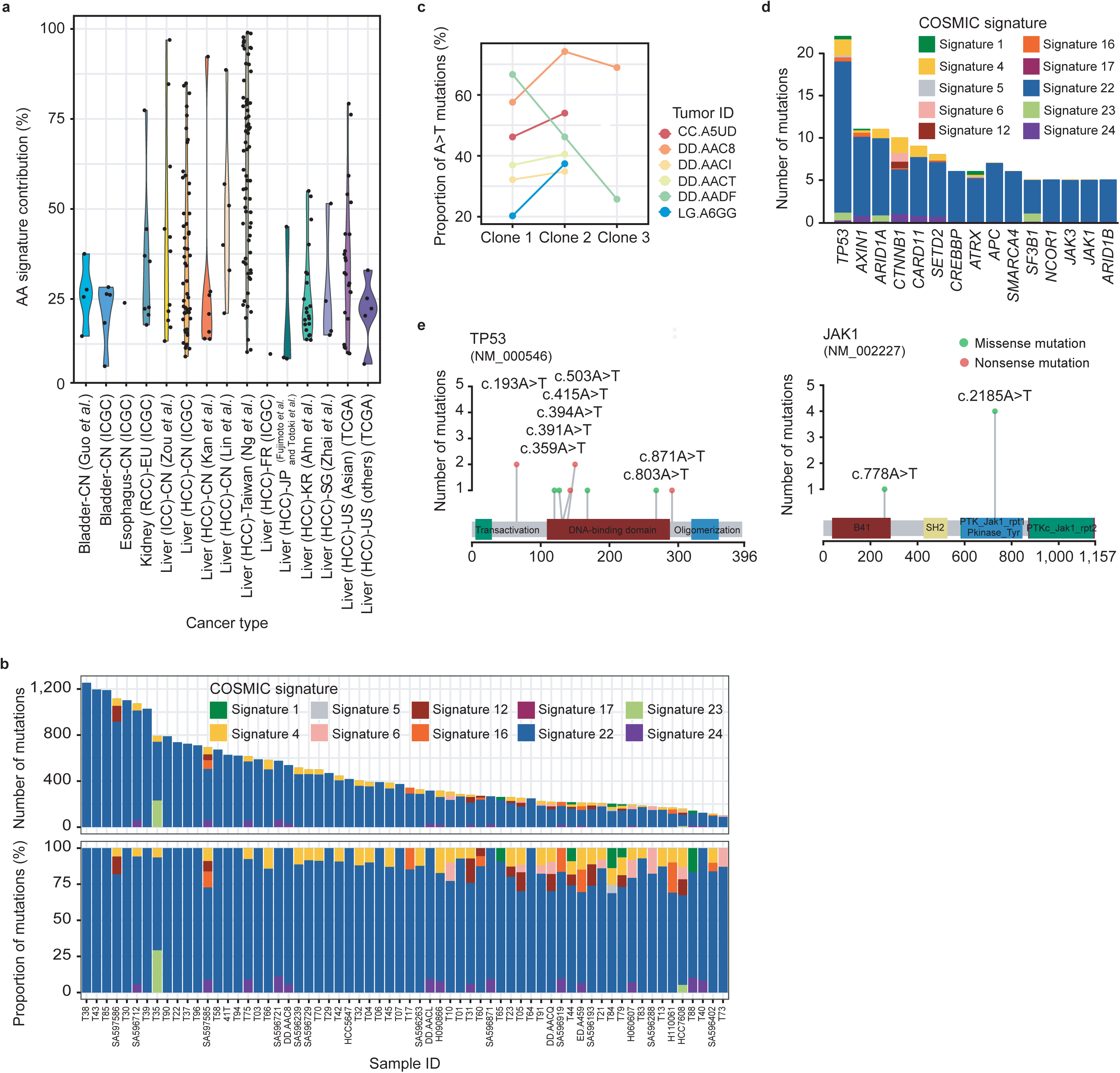
AA signatures in human liver cancer and other cancers. (**a**) The AA signature contribution in each individual of the affected human cancer types. The figure is depicted as a violin plot, in which each dot represents one human tumor. (**b**) COSMIC signature contribution according to mutational counts (upper) and proportions (bottom) in the selected human HCCs with an AA signature proportion larger than 52%. (**c**) The curves of the A>T mutation proportions in different subclones within 6 TCGA-derived human HCCs. The A>T mutations were deposited in the founding clones. Tumor DD.AAC8 presents a typical pattern, implying that AA led to tumor initiation and diminished in the later processes, as in the mouse tumors. The upward-pointing lines of the AA contributions across the clonal evolution in the other 5 tumors indicate sustained AA exposures in the patients. (**d**) Cosmic signature contributions to the essential driver genes reported in HCCs. (**e**) The AA signature caused A>T mutation sites in *TP53* and *JAK1* in the selected human HCCs with an AA signature proportion larger than 52%.

To uncover the affected driver mutation by AA, next we investigated the liver cancers that harbored a nonsilent or splicing site with AA signature mutations in known driver genes. Here, the mutations were ascribed to the signatures using a Bayesian classifier, which showed that 83 (16%) of the 510 HCCs from China had driver genes affected by the AA signature, while 16 (4%), 1 (11%), and 11 (5%) were respectively identified in the US, Singapore and Korea HCCs, as well as 4 (4%) ICCs from China (**Table 1**).

In addition, we also applied a more stringent threshold to estimate the AA contribution to human liver cancers by applying the lowest observed AA signature intensities in our experimental mouse tumors, requiring the 95% lower confident interval of the AA signature contribution to be larger than 52%, which was the lowest contribution of the bootstrapped results obtained for these mouse tumors (**Supplementary Fig. 5e**). We used this value as an indication of carcinogenic dosage in liver cancer. Our method appeared to perform well for detecting an exposure contribution above 50%, as the accuracy, specificity, sensitivity and F-measure all equaled 1 (**Supplementary Fig. 5c**). As a result, 64 (3%) of the 1957 HCCs worldwide were identified as having an AA exposure greater than 52% (**Table 1** and **Fig. 6b**). Significantly, among them, 57 (89%) of the 62 were from China. Additionally, 3 (3%) of the ICCs in China were identified with the same standard.

The finding that the AA mutational signature exists in human cancer, particular in Chinese liver cancer, based on the different criteria (**Table 1**), strongly indicates that AA exposure in Chinese population might have been one of the major risk factors for the onset of liver cancer, including HCC and ICC.

### AA exposure could be operative in an earlier stage of human liver cancer

We hope to determine whether AA is operative in the initial stage of human liver cancer, similar to its role in mouse liver cancer. Therefore, we performed a clonality analysis in the TCGA-derived HCC samples with the AA signature and DNA copy number profiles, as we did in the mouse tumors. Here, we chose 6 HCC samples with an AA signature contribution above 30% to compare their AA exposures along with clonal evolution (**Fig. 6c**). It was shown that, in all 6 tumors, AA-associated mutagenesis was operative in the initial subclones. In sample DD.AADF, the AA signature was found to be the predominant etiology in the founding subclone and decreased in later formed subclones, demonstrating a similar trend to that observed in mouse liver cancers. In contrast, the AA signature seemed to continue or even increase throughout the clonal evolution of the other five human HCCs, possibly due to the prolonged AA exposure in patients rather than acute exposure in our mouse model. In addition, among 3 of 9 Singapore patients with HCCs carrying the AA signature, we ascribed the mutations as early (trunk) and late (branch), as the study provided multisector sequencing results. It was shown that trunk mutations had higher proportions of A>T transversions, indicating that AA exposure was likely to play roles in the initial stage of tumorigenesis (**Supplementary Fig. 5f**).

Moreover, we analyzed the contribution of the AA signature to known driver genes in HCCs. Excluding *TP53, ARID2, ARID1A* and *AXIN1* frequently harbored nonsilent A>T mutations (**Fig. 6d**), which encode components of SWI/SNF complex and Wnt-β-catenin pathway that were also disrupted in AAI-induced mouse liver cancer.

In addition, to identify the driver mutations induced by AA exposure, we further investigated the cancer genomic data regarded as AA-induced liver cancer according to a more stringent criterion referring to an AA signature > 52%. We retrieved the genomic data from 62 HCCs meeting the criterion to obtain 74045 putative mutations, and we performed MutSigCV analysis, which revealed that the scattered A>T mutations in the tumor suppressor gene *TP53* were significantly affected in AA dominant human liver cancer. *TP53* (q < 0.1), along with eight A>T transversions, was significantly mutated in these AA-related liver cancers (**Fig. 6e**), where the A>T transversions led to nonsense mutations that dispute the structure of TP53, or missense mutations that alter its functions by the mutated DNA-binding domain.

We also ascribed each mutation to a specific signature and then selected the AA signature A>T mutations **(Supplementary Table 15**) for analysis via the oncodriveCLUST algorithm, a positional clustering method, because most of the oncogenic mutations of one gene were enriched at a few specific loci (aka hot-spots). Significantly, 17 genes with FDR smaller than 0.1 were identified (**Supplementary Table 16**). It is noteworthy that JAK1 S729C induced by a c. 2185 A>T mutation was identified (FDR = 5 × 10 ^-4^) as a candidate driver (**Fig. 6e** and **Supplementary Table 16**), as there were four hits at the exact same locus in *JAK1*. Interestingly, the mutation was found in Chinese liver cancer and validated as an oncogenic driver in a previous study^23^.

## Discussion

Liver cancer is the seventh most common cancer and the third leading cause of cancer-related death worldwide^29^. In China, its incidence and mortality rate are higher^30^. Liver cancer has several known risk factors, including infection with HBV and hepatitis C virus, alcohol consumption, and aflatoxin B1 contamination of food. Recently, AA has been statistically associated with human liver cancer, especially in Chinese patients^8,10^. However, no experimental evidence supports the notion that AA can directly lead the liver cancer. Significantly, our results demonstrate that AAI can directly induce mouse liver cancer, including HCC and ICC, in a dose-dependent manner, and increases the incidence of liver cancer when the liver is injured, such as CCl_4_ administration. This finding is consistent with clinical practice in China and some Asia countries, because some Chinese patients, including hepatitis patients, often take traditional Chinese herbal remedies that could contain AA. In fact, in human liver cancer, we found the characteristic AA signature in HCC and ICC patients based on very stringent criteria, indicating that AA exposure was the leading cause in some liver cancers, especially in Chinese patients. Therefore, our animal experiments and analyses of human liver cancer strongly indicate that AA can directly lead to liver cancer and should be listed as major risk factor for liver cancer.

AA exposure could be prone to trigger some driver mutations by characteristic A>T transversions, which lead to a growth advantage of the malignant transformed clones. Significantly, AAI-mediated mutations are found to be the early event during malignant clonal evolution in mouse and human liver cancer. AAI-DNA adducts could be detected not only in livers from mice exposed to AA but also in multiple heterogeneous subclones within the same tumor nodules, and the AAI-mediated characteristic A>T mutational signature was found in the founding subclones in both mouse and human liver cancers, further supporting the critical nature of AA exposure in some forms of liver cancer.

AA could prefer to damage different driver genes in different species, although these mutations share similar A>T transversions. A previous study had shown that the *ras* family, including *Hras, Kras* and *Nras*, had the same activating mutation — Q61L (CAA to CTA) — in oral administration AA-induced rat tumors^14^. In this study, the same activating mutation in *Hras* and *Kras* also occurred in most AA-induced HCC samples (*Hras*, 8/11; *Kras*, 1/11), indicating that the Ras pathway is crucial in AAI-induced mouse liver cancer. Interestingly, activating mutations of *KRAS* are frequent in human ICC^31^, although relatively lower in human HCC.

More interestingly, AA-mediated mutations also alter other genes that can lead to the deregulation of some signaling pathways, such PI3K-AKT, the chromatin remodeling SWI/SNF complex, epigenetic regulation, and the development-related Hippo, Notch and Wnt pathways (**Figure 4**), which are often associated with human HCC and ICC. Like human HCC, AAI-induced mouse HCCs also express hepatic stem or progenitor cell-related biomarkers such as *Afp, Gpc3, Dlk1* and *Prom1* (**Supplementary Fig. 3k**). In addition, it should be pointed out that, although *Tp53* and *Jak1* mutations were not found in the AAI-induced mouse HCCs, some point mutations of *TP53* and *JAK1*, especially *JAK1* S729C, could be considered as candidate biomarkers for AA exposure, similar to TP53 R249S for aflatoxin B1 contamination.

In conclusion, this study provides documented evidence indicating that AA can directly induce mouse liver cancers, including HCC and ICC, similar to the genetic pathogenesis of human liver cancers. In light of the animal model, AA exposure is considered as a major risk factor for some human liver cancers, especially among Chinese patients. The featured mutational process during malignant clonal evolution in AA-induced liver cancer reveals that AAI-mediated characteristic mutations are the earliest genetic event in tumorigenesis. Our data lay a solid foundation for the prevention and diagnosis of AA-associated human cancers, especially liver cancer.

### URLs

ICGC data portal, https://dcc.icgc.org/; TCGA data portal, https://portal.gdc.cancer.gov/; COSMIC mutation signatures, http://cancer.sanger.ac.uk/cosmic/signatures; COSMIC cancer census genes, http://cancer.sanger.ac.uk/census; HMMcopy (v1.22.0), http://www.bioconductor.org/packages/release/bioc/html/HMMcopy.html; MsigDB, http://software.broadinstitute.org/gsea/msigdb; R package pracma (v2.1.4), https://cran.r-project.org/web/packages/pracma/index.html; mSignatureDB, http://tardis.cgu.edu.tw/msignaturedb/Browse/.

## Methods

### Mice

The mice used in this study were on a C57BL6/J background. The wild-type C57BL6/J mice were purchased from the Slaccas Company (Shanghai, China). LoxP-flanked (floxed [f]) *Pten* (*Pten*^*f/f*^) mice (The Jackson Laboratory) and Alb-Cre mice (The Jackson Laboratory) were crossed to generate conditional liver-specific PTEN-KO mice designated as *Pten*^*LK*O^. All animal experiments were conducted under procedural guidelines and severity protocols with the approval granted by the Institutional Review Board on Bioethics of Shanghai Jiao Tong University. **HCC induction.** As mentioned in the results section, the male mice were randomly grouped and administered with AAI (2.5 or 5 mg/kg, dissolved in PBS) or a combination of AAI (2.5 mg/kg) and CCl_4_ (0.5 ml/kg, dissolved in corn oil) by intraperitoneal injection for different doses and times since the age of 1 or 2 weeks. Additional groups were injected with CCl_4_ alone or vehicle as controls. The exposure timeline was presented in **Fig. 1a**. Mice were sacrificed with CO_2_ anesthesia. The visible discrete tumors at mice livers were dissected and counted. Tumor sizes was measured with a caliper at its largest diameter. No mice were excluded in subsequent analyses.

### Histology, immunohistochemistry and immunofluorescence

Formalin-fixed tissues were embedded in paraffin. Sections (5 μm) were stained with hematoxylin and eosin (H&E), and PicroSirius Red. For immunohistochemical (IHC) staining, sections were incubated with primary antibodies against AFP (polyclonal rabbit, 1:100; ab46799, Abcam), Ki67 (polyclonal rabbit, 1:200; ab15580, Abcam) and CK19 (monoclonal rabbit, 1:400; ab52625, Abcam) overnight at 4 °C. HRP - conjugated anti-rabbit secondary antibody (polyclonal goat, 1:400; A0545, Sigma) and DAB (Sangon Biotech, Shanghai, China) were used to detect the primary antibodies, followed by hematoxylin redyeing. For immunofluorescence assay, tissues were embedded in OCT (optimal-cutting-temperature compound). Cryosections (5 μm) were fixed in 4% paraformaldehyde for 10 min and then permeabilized with 0.1% Triton X-100. Sections were incubated with primary antibody against γ-H2AX (monoclonal rabbit, 1:200; 9718, Cell Signaling) overnight at 4°C. Primary antibody were detected using fluorescent-conjugated secondary antibody (polyclonal Donkey, 1:1000; A21206, Invitrogen). The nuclei were stained with DAPI and mounted with anti-fading mounting reagent (Fluoromount™ Aqueous Mounting Medium, Sigma). Brightfield images were taken using a Nikon Eclipse Ni microscope. Fluorescence images were taken using fluorescent confocal microscope (Nikon A1Si). Sirius Red stained area or fluorescence intensities of 10 non-overlapping fields in each section were quantified using Fiji Image J at × 100 or × 400 magnification.

### Western blot analysis

Liver tissues were lysed and the protein concentrations were determined using the BCA assay (Thermo Scientific). Membranes were incubated with the following primary antibodies: anti-γ-H2AX (monoclonal rabbit, 1:1000; 9718, Cell Signaling), anti-p53 (monoclonal mouse, 1:200; sc-126, Santa Cruz), anti-Bax (polyclonal rabbit, 1:200; sc-493, Santa Cruz), anti-p-ATR (Ser428, polyclonal rabbit, 1:1000; 2853, Cell Signaling), anti-AFP (polyclonal rabbit, 1:1000; ab46799, Abcam), GPC3 (polyclonal rabbit, 1:400; ab66596, Abcam), E-cadherin (polyclonal rabbit, 1:1000; 20874-1-AP, Proteintech), p-ERK (Tyr 204, monoclonal mouse, 1:200, sc-7383, Santa Cruz), ERK1 (polyclonal rabbit, 1:400, sc-94, Santa Cruz), p-AKT (Ser473, monoclonal rabbit, 1:2000; 4060, Cell Signaling), AKT (polyclonal rabbit, 1:1000; 9272, Cell Signaling) and YAP (polyclonal rabbit, 1:1000; 4912, Cell Signaling) and anti-β-actin (monoclonal mouse, 1:5000; A2228, Sigma). HRP-conjugated secondary antibodies (polyclonal goat, 1:10000; A6154 and A4416, Sigma) were applied. Proteins was detected by ECL reagent (Share-bio, Shanghai, China).

### In vivo alkaline comet assays

The male mice (n = 4) were administered with PBS (10 ml/kg) or AAI (2.5 or 5 mg/kg) by intraperitoneal injection at the age of 2 weeks. Mice were anesthetized at 3 hours after administration. Livers were perfused with Hanks’ balanced salt solution (HBSS). Then alkaline comet assay was performed with CometAssay kit (4250-050-K, Trevigen) following the manufacturer’s instructions. Afterwards, the slides were stained with SYBR Green I (Sangon Biotech, Shanghai, China). Fluorescence images were taken using fluorescent confocal microscope (Nikon A1Si) at × 100 or × 400 magnification. Tail DNA were analyzed using the CASP^32^. At least 100 cells were randomly selected and analyzed per sample. A total of 600 cells were analyzed per group.

### DNA and RNA extraction

Tissues were minced and digested overnight at 55 °C in 10 mM Tris -HCl (pH 8.0) containing 100 mM EDTA, 10 mM NaCl, 0.1% SDS, proteinase K (0.2 mg/ml), and RNase A (0.2 mg/ml). DNA was purified by phenol/CHCl_3_. RNAwas extracted from frozen tissue using TRIzol according to the manufacturer’s instructions.

### Synthesis and identification of dA-AL-I

7-(deoxyadenosin-N6-yl) aristolactam I (dA-AL-I) was synthesized by incubating deoxyadenosine (dA) with AAI according to the method described previously^33^. A mixture containing dA-AL-I, AAI and dA was yielded.

Synthetic dA-AL-I was analyzed with an ACQUITY ultra performance liquid chromatography (UPLC) system (Waters) connected to a XEVO-G2XS quadrupole time-of-light (QTOF) mass spectrometer (UPLC-QTOF-MS) (Waters) with electron spray ionization (ESI). Seven microliters synthetic dA-AL-I was injected into an ACQUITY HSS T3 column (2.1 mm × 100 mm i.d., 1.8 μm particle size) (Waters) in positive electrospray ionization mode at a flow rate of 0.4 ml/min. Mobile phase A and B were 0.1% formic acid in water and acetonitrile, respectively. The gradient program used was: 0-1 min, 1% B; 1-3 min, 1-30% B; 3-7 min, 30% B; 7-9 min, 30-100% B; 9-11.2 min, 100% B; 11.2-11.3 min, 100-1% B and 11.3-13 min, 1% B. The ESI source was operated in positive ion mode with a capillary voltage of 2 kV, cone voltage of 40 V, source temperature of 115 °C, desolvation temperature of 450 °C, cone gas flow of 50 l/h, and desolvation gas flow of 900 l/h. The mass spectra were acquired over m/z 50-1200 in full scan mode. The secondary mass spectra (MS/MS) were also acquired in the positive mode in the range of m/z 50-600 with the collision energy of 10-30 eV. Data were acquired by software Masslynx v 4.1 and analyzed by UNIFI 1.8.1.

### Mass spectrometry identification and quantitation of AAI-DNA adducts

Liver and renal tissue DNA (500 μg and 50 μg in 5 mM bis-tris-HCl buffer (pH 7.1) containing 10 mM MgCl_2_) was digested with DNase I (Worthington), nuclease P_1_ (Sigma), alkaline phosphatase (Worthington) and phosphodiesterase I (Worthington) as described previously^34^. Protein was precipitated by adding 2 vol of chilled C_2_H_5_OH and centrifuging. The supernatant was concentrated by vacuum centrifugation and dissolved in a solvent of 1:1 H_2_O/DMSO (50 μl)^34^.

Ultra-high-performance liquid chromatography/triple quadrupole mass spectrometry (UHPLC/QQQ MS) was used for identification and quantitation of AAI-DNA adducts. The chromatographic column and mobile phases used with this system were the same as those that were used with the UPLC-QTOF-MS system. The injection volume of synthetic dA-AL-I or DNA digestion product was two microliters. The QQQ MS system (Waters) was operated in positive ion mode with a capillary voltage of 0.5 kV, cone voltage of 30 V, source temperature of 150 °C, desolvation temperature of 500 °C, cone gas flow of 150 l/h, and desolvation gas flow of 1000 l/h. Detection of the ion pairs were performed by multiple reaction monitoring (MRM) mode. The MRM ion pairs for dA-AL-I were 543.16/427.12, 543.16/395, and 543.16/292, and the corresponding collision energies were 25, 30 and 35 eV, respectively. The quadrupoles were set at unit resolution. The analytical data was processed by Masslynx.

### WGS and copy number analysis

Whole-genome DNA libraries were created with Illumina Truseq Nano DNA HT Sample Prep Kit following the manufacturer’s instructions. The libraries were sequenced on Illumina Hiseq platform and 150 bp paired-end reads were generated. We investigated the copy number patterns of the samples applying the suite of HMMcopy (v1.22.0) on the WGS data. Briefly, the coverages were initially corrected with the GC and mappability bias of the reference genome. Then the corrected signals were segmented using Hidden Markov Model to yield an estimate of the copy number events.

### WES and somatic mutation calling

Whole-exome capture was done with Agilent SureSelect Mouse All Exon V1 kit according to the manufacturer’s instructions. The libraries were sequenced on Illumina Hiseq platform and 150 bp paired-end reads were generated. Raw sequencing reads of WES were aligned against mouse reference build GRCm38 using bwa (v0.7.11)^35^. The duplicates were removed by Picard Tools (v1.4.5) (http://broadinstitute.github.io/picard/). The base quality was recalibrated using the Genome Analysis Toolkit (GATK 3.7-0-gcfedb67)^36^. Mutect (v2.0)^37^ was employed to predict somatic mutations of liver tumor and adjacent tissue with the corresponding tail tissue being the control. The mutations were removed if it was in mouse dbsnp. Filtered somatic mutations were functionally annotated by ANNOVAR^38^, using the RefGene database. Nonsynonymous, stop-loss, stop-gain and splice-site SNVs (based on RefGene annotations) were considered to be functional. SNPEFF (v4.3s)^39^ were used to predict functional influences of the somatic mutations. Bam files were visualized in Integrated Genome Viewer (IGV)^40^. Nonsynonymous mutant genes in AAI-induced mouse liver cancer were performed with KEGG pathway enrichment analysis^41^.

### RNA sequencing, analysis and annotation

RNA-seq libraries were generated using NEBNext^®^ Ultra™ RNA Library Prep Kit for Illumina^®^ according to the manufacturer’s instructions. Then they were sequenced on an Illumina Hiseq platform to generate 150 bp paired-end reads. Sequenced reads were mapped to the GRCm38 UCSC annotated transcripts via Tophat (v2.1.0)^42^. Transcripts were then assembled and counted with the Cufflinks suit (v2.2.1)^43^. Differentiated expressed genes were analyzed by Cuffdiff.

### Gene set enrichment analysis

Gene set enrichment analysis (GSEA v3.0)^44^ was performed using the normalized expression values generated by Cuffnorm between the liver and the tumors. Differential enrichment was calculated using the signal-to-noise metric. FDR 0.1 was set as significant in the analysis. To investigate the response of expression alterations to the significantly mutated pathways, we investigated the gene sets of the target genes of the activated transcription factors, such as Ets1 in the Ras pathway. Therefore, the analysis was run using the ‘motif’^45^, together with ‘KEGG’, and ‘GO’ signature collections from the Molecular Signature Database (MsigDB). Differentially expressed genes in the Ras, Hippo and PI3K-AKT downstream transcription factor associated gene sets (FDR < 0.1) were selected and listed in **Supplementary Table 11**, along with differentially expressed the liver cell stem markers^46^, and Wnt (http://web.stanford.edu/group/nusselab/cgi-bin/wnt/) and Notch^47^ signaling pathway target genes. The top 30 genes with higher fold change were used to generate the heatmap in **Supplementary Fig. 3k**.

### Clonal and phylogenetic reconstruction

With the information of copy number and mutation allele frequency, Sciclone was used to characterize coexisting subpopulations in the individual tumors, both in the mouse liver tumors and the TCGA-derived liver cancers. The minimum depth of coverage was set as 70-fold and 50-fold respectively for mouse and human data. The phylogenetic tree of the 11 tumor nodules was reconstructed via R package ape (v5.2)^48^ with the application of the neighbor-joining algorithm^49^. R package fishplot^50^ was used to visualizing tumor evolutions of the discrete tumor nodules within the same mouse.

### Mutation signature analysis

Trinucleotide contextualized mutational signature deconvolution was previously described as cocktail party problem^51^. We used the least square root implemented in the R package pracma to decipher the mutational signatures with the known mutational signatures inferred in the specific cancer type, from “Signatures of Mutational Processes in Human Cancer” in the COSMIC database, as recommended in the signature analyzing R package Mutational Patterns (v1.8.0)^52^. Briefly, the algorithm deciphers the set of mutational signatures that optimally explains the total trinucleotide frequencies. For the mice tumor signature deciphering, we adopted this method with a cutoff of 5% of signature contribution to avoid over fitting

For human cancer signature investigation, we initially used the webserver mSignatureDB^53^ to investigate the COSMIC signature contributions across 73 research programs over 15, 780 tumors documented on The Cancer Genome Atlas (TCGA) and the International Cancer Genome Consortium (ICGC) data portals using the deciphering method provided on the server. Next, we used the locally adopted least square root implementation to double check the positively detected cancer projects. To improve the deconvolution, bootstrap resampling implemented in the R package Signature Estimation^54^ was employed to calculate the confident interval of signature exposures. In this way, we conducted 1000 times of randomized re-sampling in order to simulate the perturbation of the input data. Then we generated estimation of the exposures of the mutational signatures in each bootstrap sample. From the continuum of the estimated signature contributions, we retrieved the lower boundary of the 95% confidence intervals of the bootstrapped AA signature distribution to obtain a probability of 0.05 for rejection of the event that the AA signature contribution being above the specific retrieved threshold. To evaluate the performance of the deciphering procedure, we used the random sampling and permutation function to generate a simulated dataset of 1000 samples with known signature exposures and mutation counts. 156 samples were excluded for containing zero mutation. The remained 864 samples were used for evaluation of deciphering methods. There was a work reporting that performing the same method on the exome resided mutations or the genome resided mutations revealed different results; the former was stricter for AA signature detection^55^. It is to some degree due to the overfitting danger when dealing with a large load of mutations. Therefore, to avoid sequence bias between WES and WGS generated data, we only retained the exome resided mutations from WGS generated data for the mutation deconvolution analysis.

To estimate similarities between tumor or clonal mutational profiles, and the COSMIC signatures, cosine similarities were calculated using the R package Mutational Patterns (v1.8.0).

### A>T Transcriptional strand bias

For each tumor, A>T transcriptional strand bias was analyzed by comparing the number of mutations occurring on the transcribing and non-transcribing strands over the genome with the Poisson distribution test. Later on, to correlate AAI mutational processes with gene transcription history, we categorized the UCSC (University of California Santa Cruz)^56^ known genes into 5 categories from no to high transcriptional activities in the RNA-seq data of the 11 non-tumor liver samples and compared the transcriptional strand bias within each defined gene expression category.

### Mutation assignment to the signatures

Each mutation was firstly ascribed to a specific signature via a Bayesian inference method implemented in the R package Palimpsest ^57^, which was calculating the probability of each operative process for a certain mutation and then choosing the largest as the assigned signature for the specific mutation.

### Driver gene analysis

To analyze the significant A>T mutated genes in the mice, we calculated the nonsilent mutational counts per mega base for each gene to search for the genes that are mutated more frequently. The genes listed in the duplicated gene database were removed as they are easily to be falsely detected with mutations^58^. The MutSigCV^59^ and oncodriveCLUST^60^ analyses were performed on the human mutation data for drive gene identification. R package maftools^61^ was used to plot the gene mutation points distribution on the motifs and Palimpsest was used to calculate the contribution of the operative signatures to the reported driver genes in HCC^57^.

### Statistics

Statistical analyses were performed using SPSS software. All the statistical tests used were described in the relevant sections of the manuscript. *P*-values < 0.05 were considered statistically significant.

## Data availability

The mouse next-generation sequencing data used in the manuscript can be downloaded from the database of NCBI under accession number: PRJNA 507339.

## Acknowledgements

We sincerely thank associate professors Lan Wang and Kunyan He of Shanghai Center of Systems Biomedicine, Shanghai Jiao Tong University, for providing critical comments in this study. This work was supported in part by China National Science and Technology Major Project for Prevention and Treatment of Infectious Diseases (grant no. 2017ZX10203207), National Program on Key Research Project of China (grant number 2016YFC0902701) and National Natural Science Foundation of China (81472621 and 81672772).

## Author contributions

Z.-G.H. initiated and supervised the project. Z.-N.L., L.-N.Z. and X.-B.S. performed animal test, other experiments and statistical analysis. Q.L. and Y.S. analyzed the mouse WGS and WES data. Q.L. did the other bioinformatics analysis. Z.-G.H., Z.-N.L, and Q.L. analyzed the data and wrote the manuscript.

## Supplementary information

Supplementary Figures 1–5 and Supplementary Tables 1–16

