## Supplemental Figures 1-5 for "The mutational features of aristolochic acid-induced mouse and human liver cancers"

**Supplementary Figure 1. AAI can induce liver cancer in wild-type and liver-specific Pten-deficient C57BL/6 male mice.**

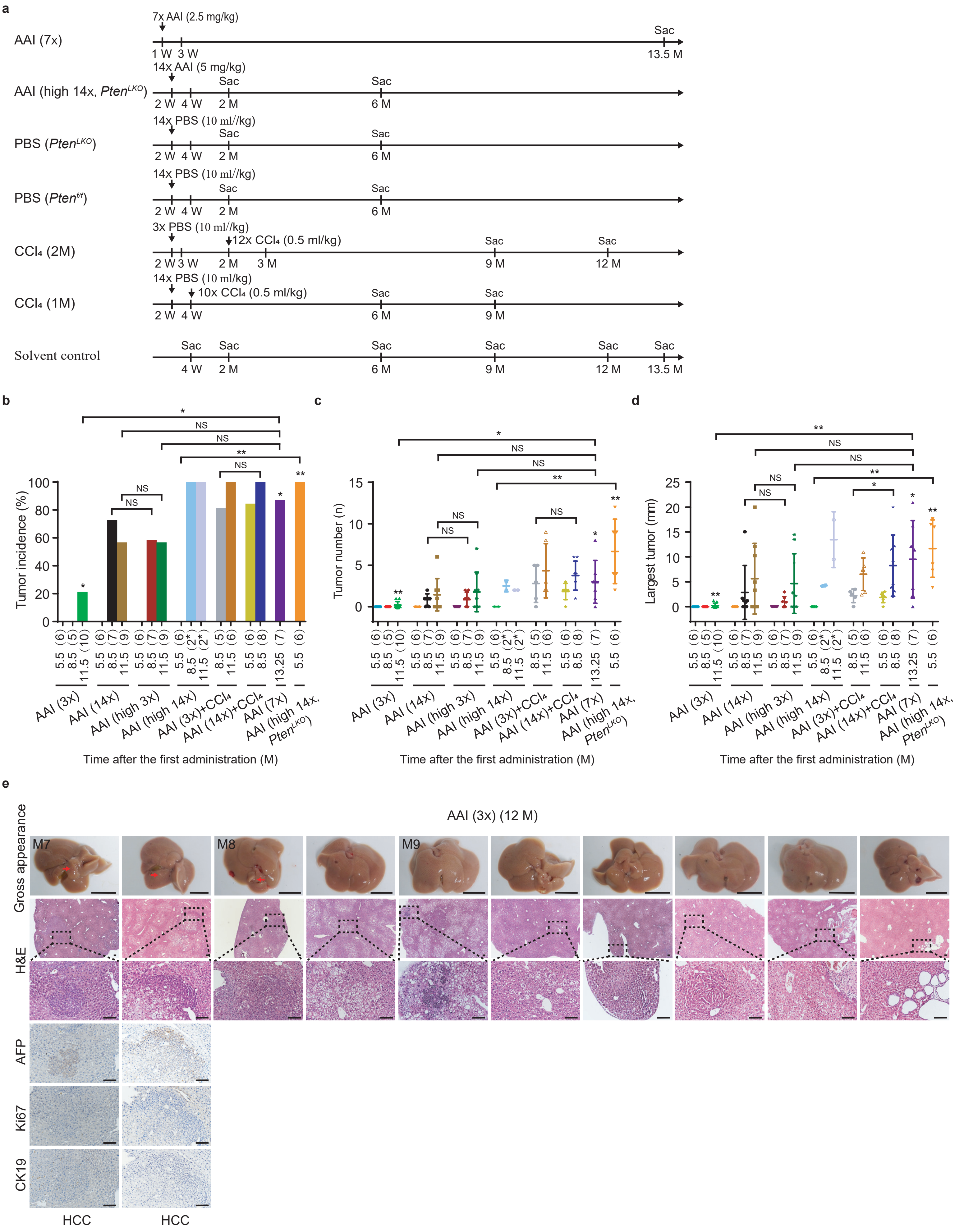

f

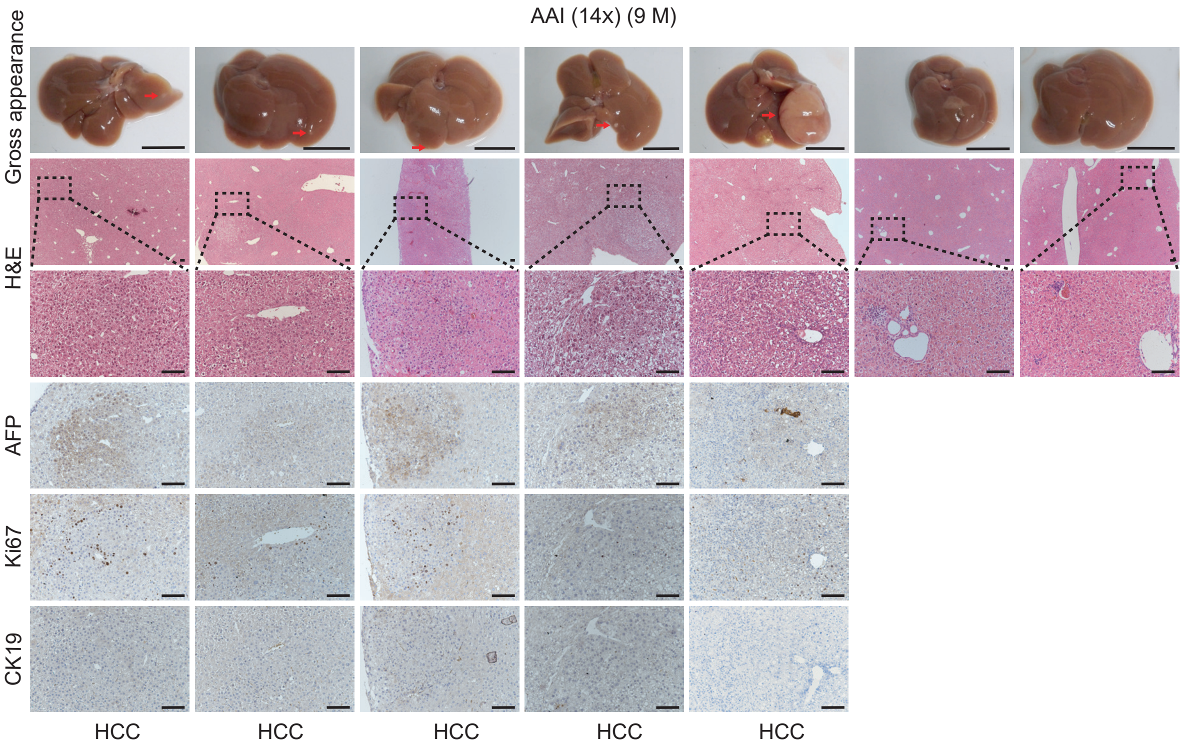
AAI (14x) (9 M)
Gross appearance
H&E
AFP
Ki67
CK19
HCC
HCC
HCC
HCC
HCC

g

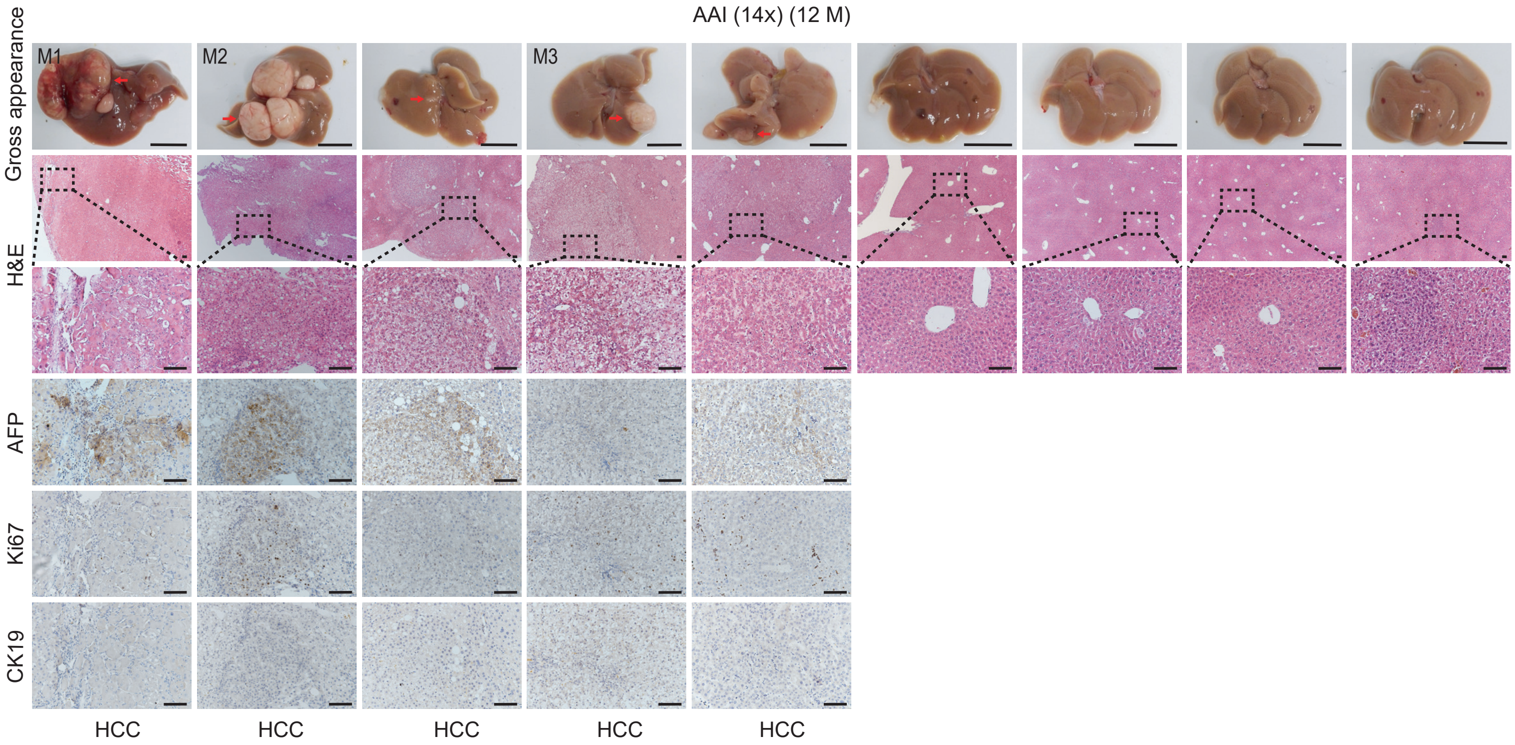
AAI (14x) (12 M)
Gross appearance
M1
M2
M3
H&E
AFP
Ki67
CK19
HCC
HCC
HCC
HCC
HCC

**h**

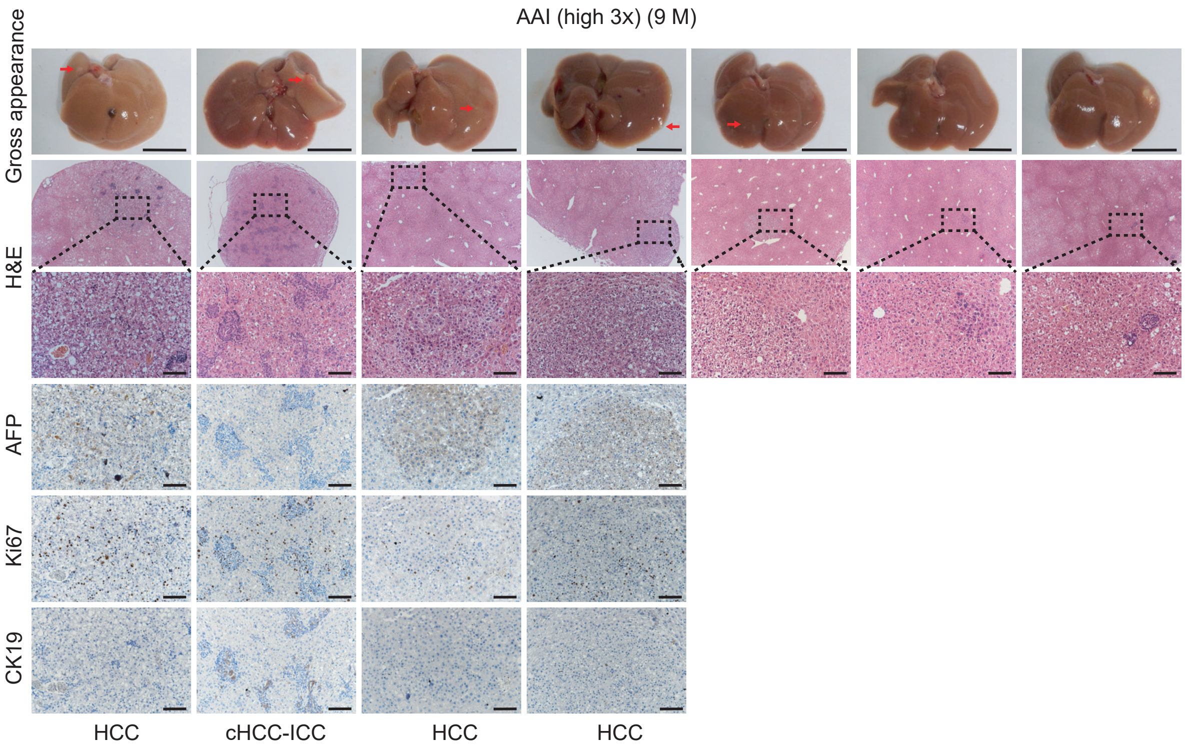

AAI (high 3x) (9 M)
Gross appearance
H&E
AFP
Ki67
CK19
HCC
cHCC-ICC
HCC
HCC


**i**

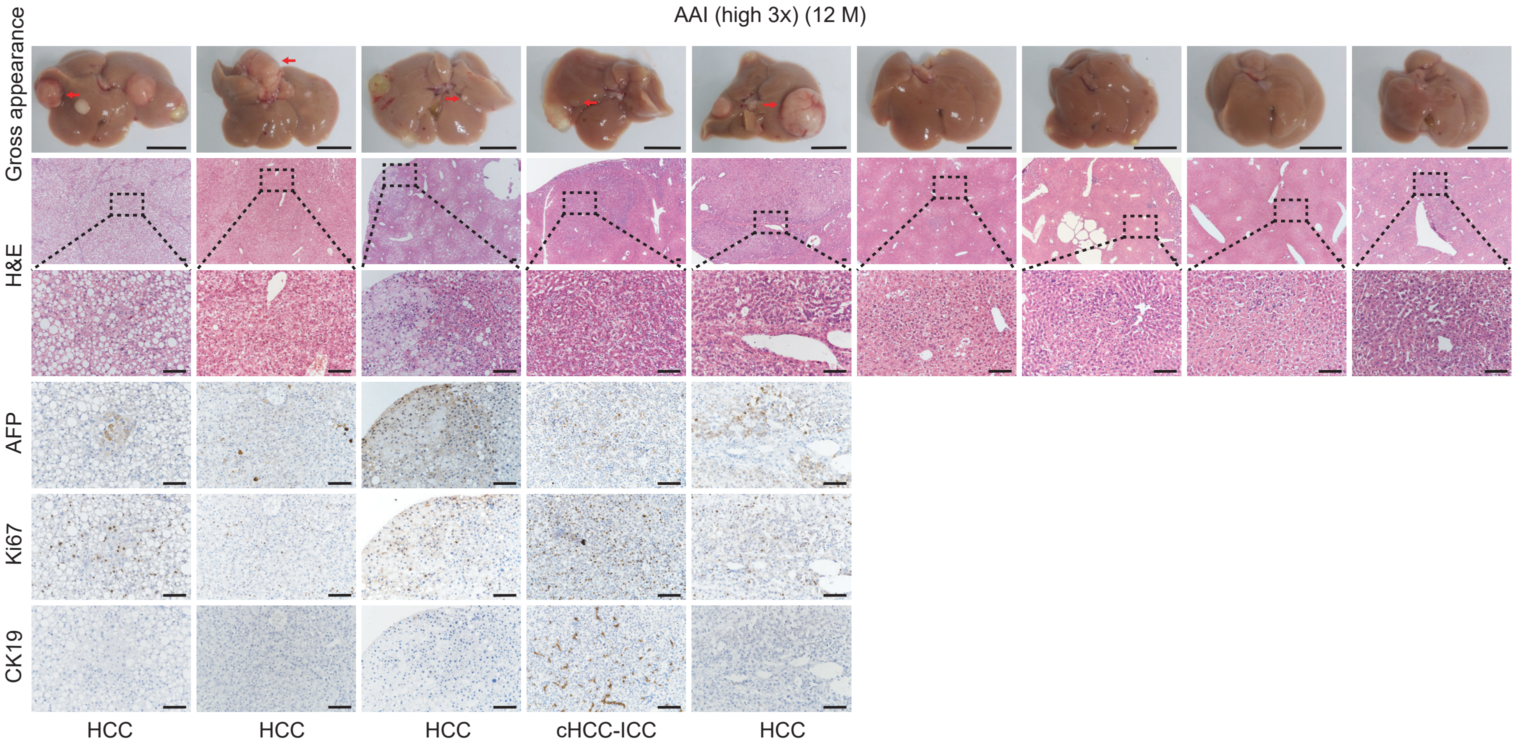

AAI (high 3x) (12 M)
Gross appearance
H&E
AFP
Ki67
CK19
HCC
HCC
HCC
cHCC-ICC
HCC

**j**

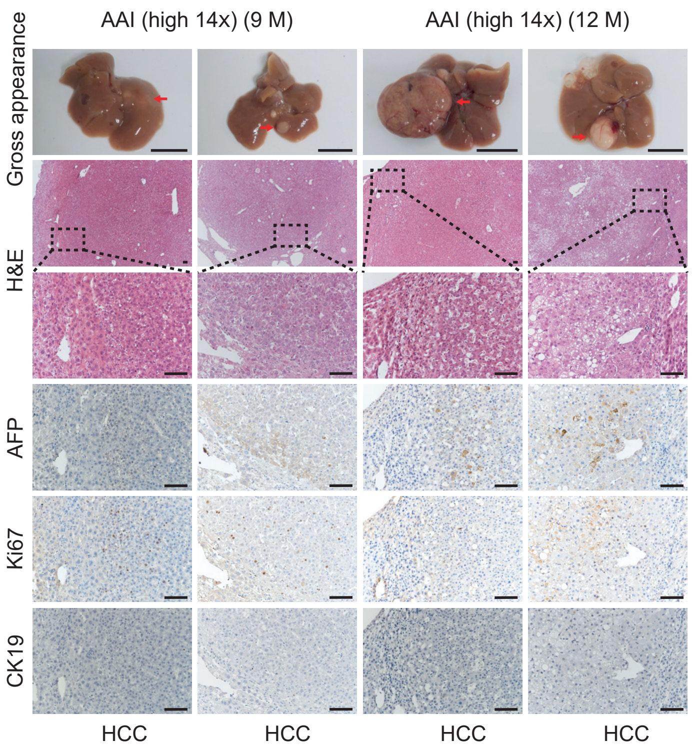
AAI (high 14x) (9 M)
AAI (high 14x) (12 M)
Gross appearance
H&E
AFP
Ki67
CK19
HCC
HCC
HCC
HCC

**k**

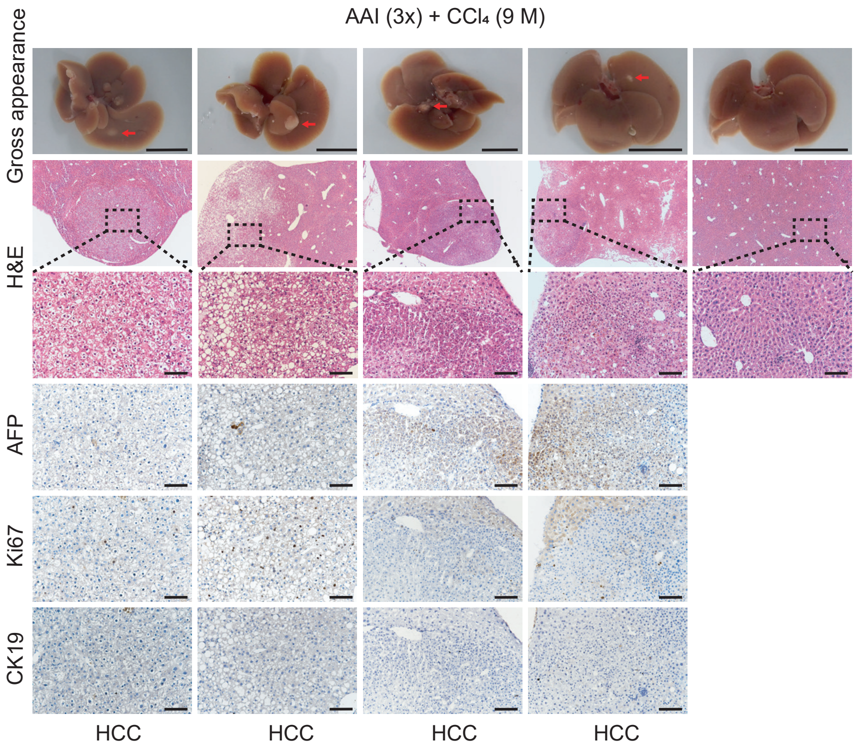
AAI (3x) + CCl₄ (9 M)
Gross appearance
H&E
AFP
Ki67
CK19
HCC
HCC
HCC
HCC

**l**

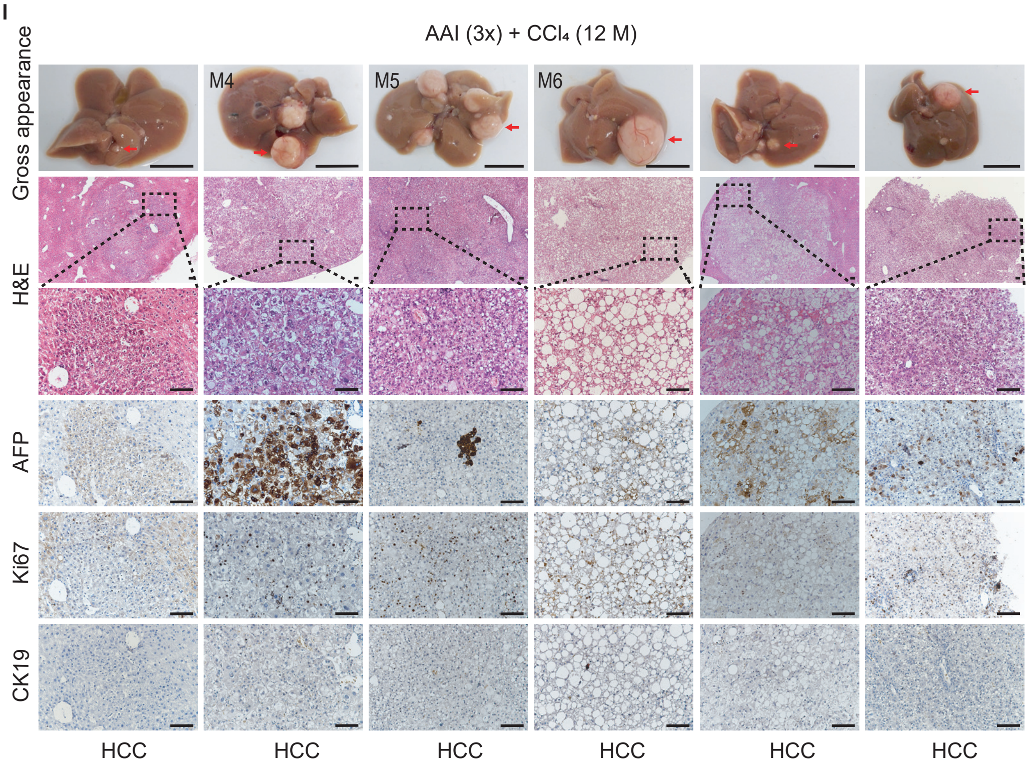
AAI (3x) + CCl4 (12 M)
Gross appearance
H&E
AFP
Ki67
CK19
M4
M5
M6
HCC
HCC
HCC
HCC
HCC
HCC

**m**

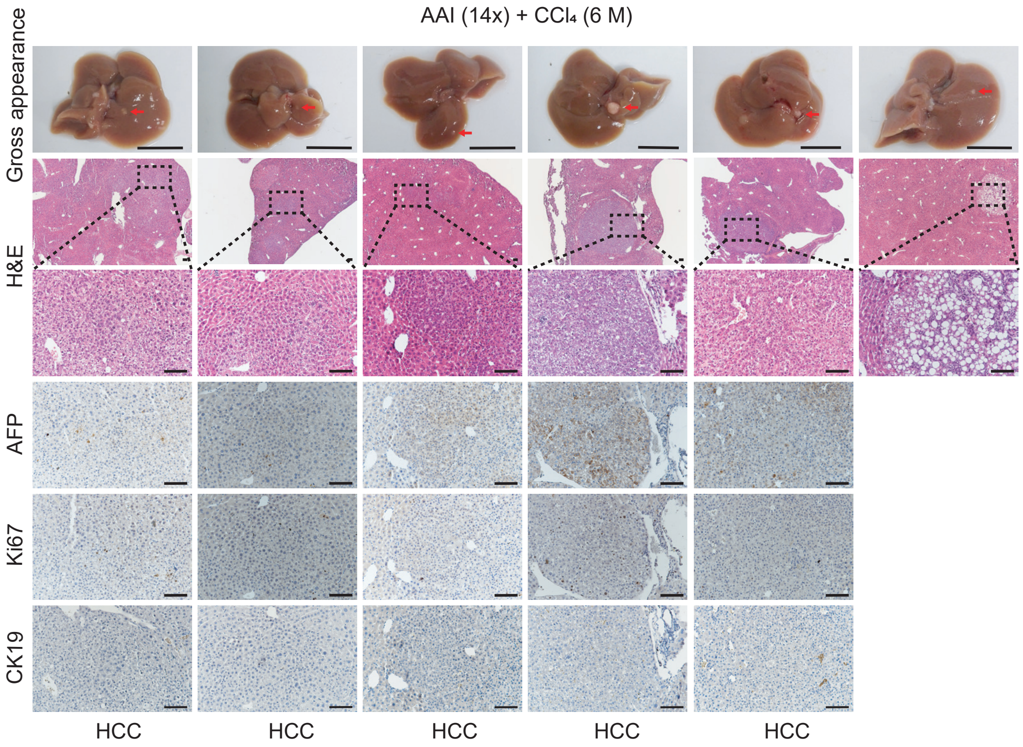
AAI (14x) + CCl4 (6 M)
Gross appearance
H&E
AFP
Ki67
CK19
HCC
HCC
HCC
HCC
HCC

n

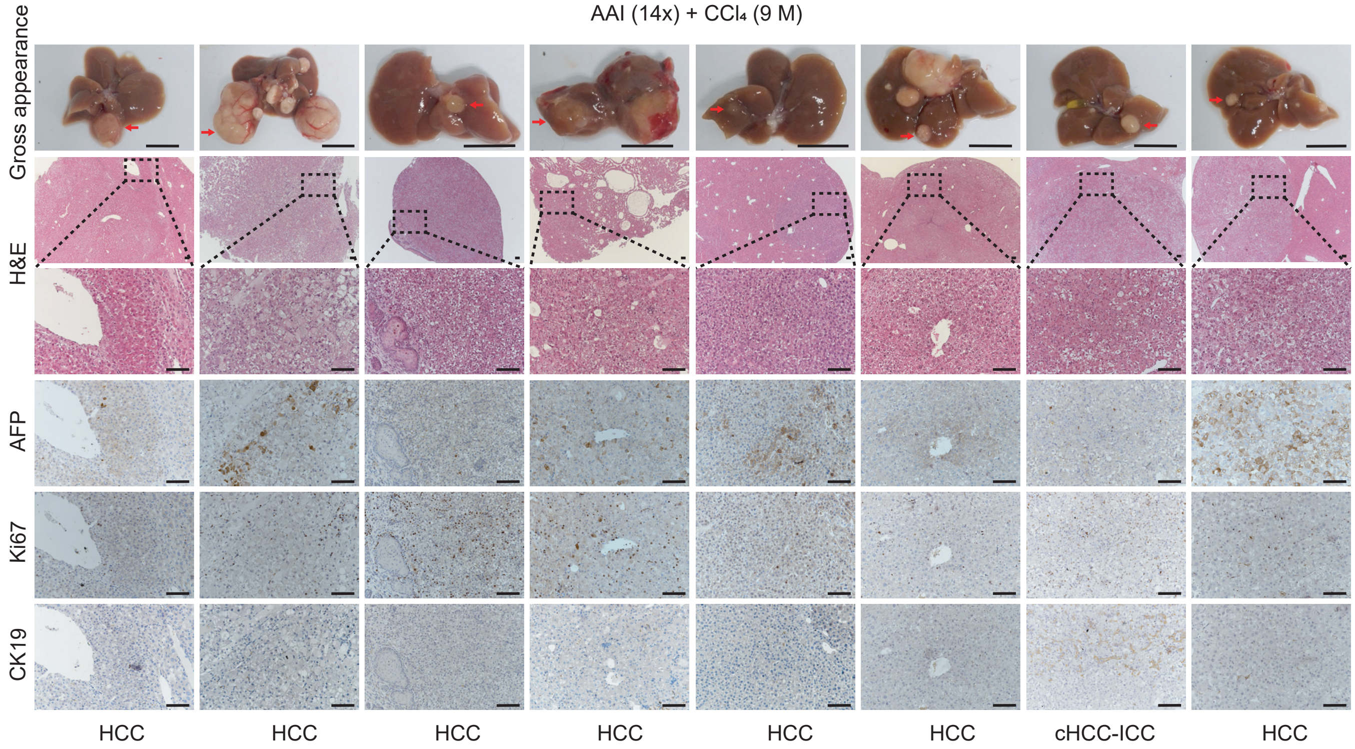
AAI (14x) + CCl4 (9 M)
Gross appearance
H&E
AFP
Ki67
CK19
HCC
HCC
HCC
HCC
HCC
HCC
cHCC-ICC
HCC

o

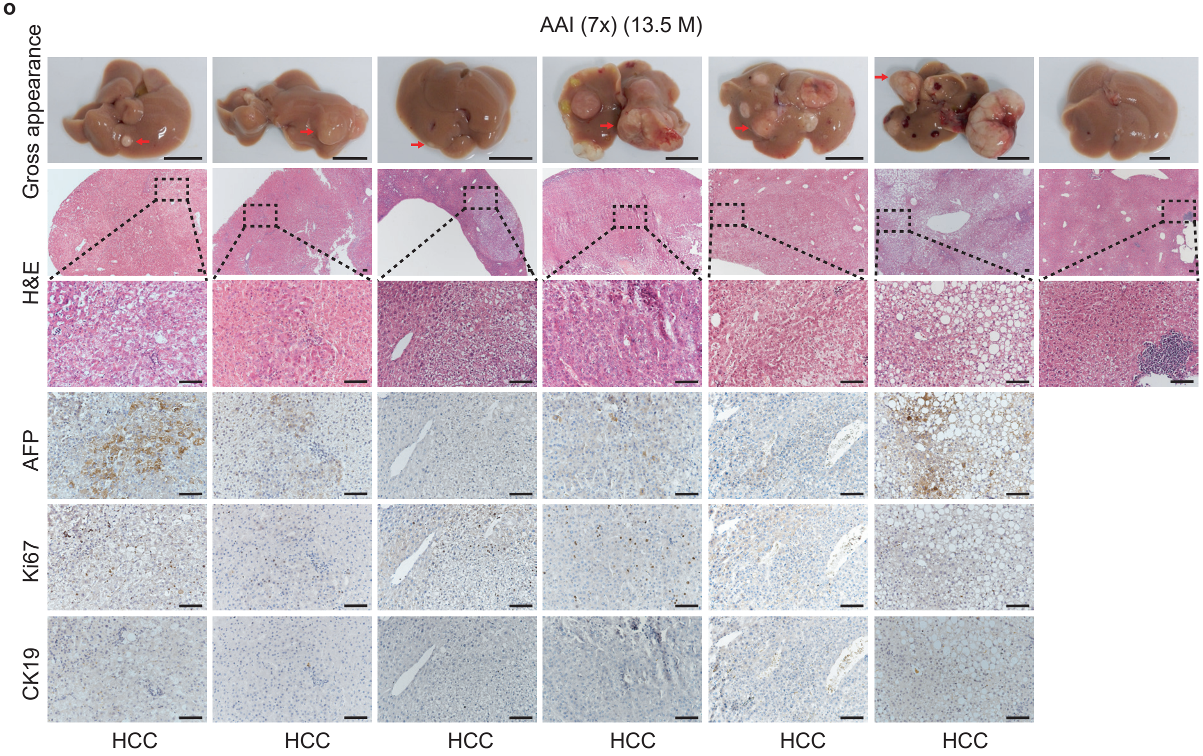
AAI (7x) (13.5 M)
Gross appearance
H&E
AFP
Ki67
CK19
HCC
HCC
HCC
HCC
HCC
HCC

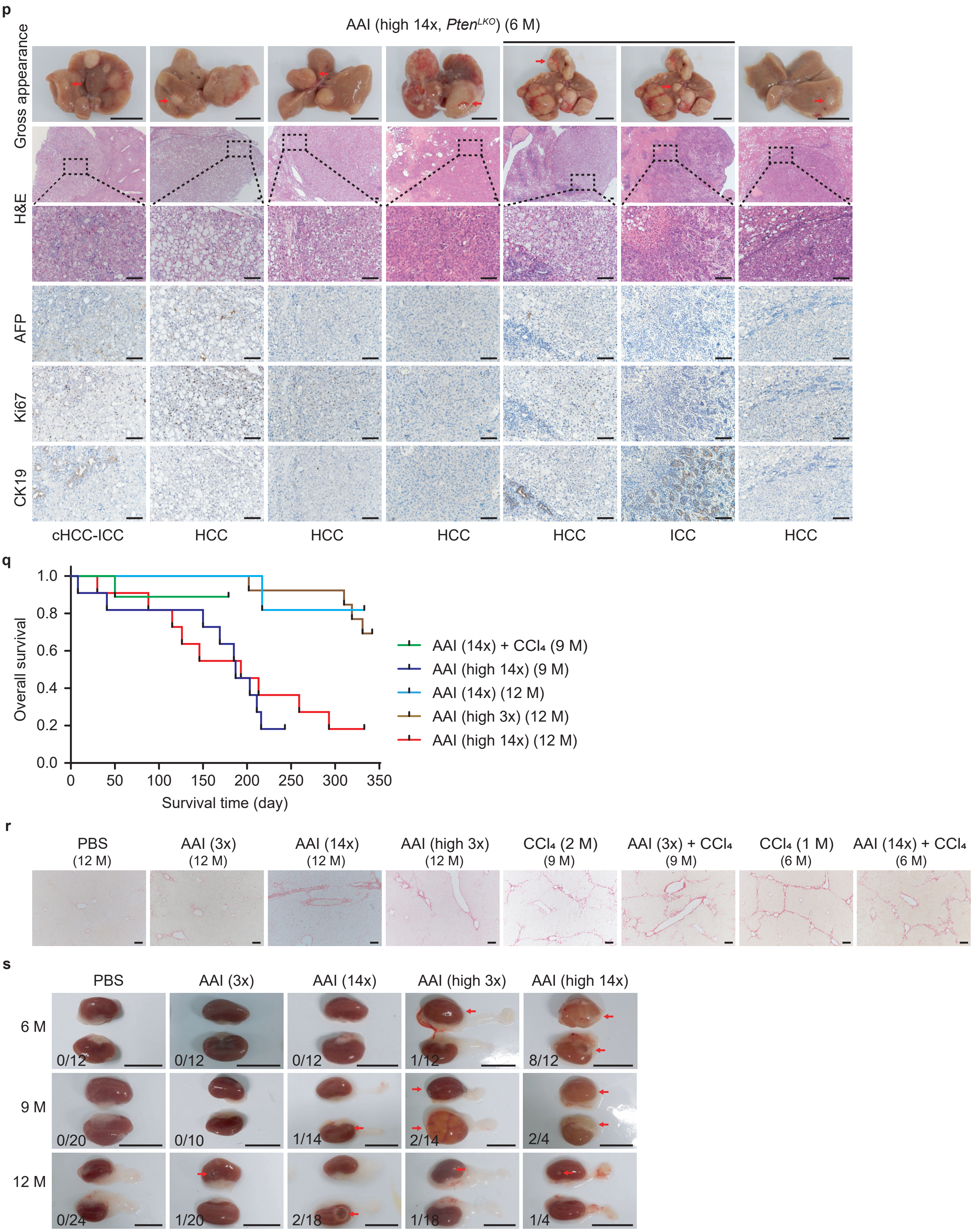


(**a**) Simplified diagram of liver cancer induction in wild-type and liver-specific Pten-deficient C57BL/6 male mice with AAI alone or a combination of AAI and $CCl_4$, where the dosages and time points of drug administration are indicated by arrows, and samples were harvested at the indicated time of mouse sacrifice (Sac). (**b-d**) Tumor incidence (**b**), tumor number (**c**), and largest tumor size (**d**) of AAI-induced liver cancer. The number after the month is the number of mice in each group. In this figure, the group "AAI (3x)" (11.5 M) was compared with 39 normal mice at 12 months to 19 months of age. The numbers of mice in the control groups corresponding to "AAI (7x)" and "AAI (high 14x, *Pten*$^{LKO}$)" in the figure were 5 and 6, respectively. The numbers of mice in the control groups correspond to other experimental groups in the figure, as mentioned in **Fig. 1**. (**b-d**) Values indicated by long and short horizontal lines represent the mean ± SD. (**e-p**) Gross appearance (scale bars, 1 cm), H&E staining (scale bars, 100 μm), and IHC analysis with anti-AFP, Ki67 and CK19 antibodies (scale bars, 100 μm) in all models of administration, including "AAI (3x)" (12 M) (**e**), "AAI (14x)" (9 M) (**f**), "AAI (14x)" (12 M) (**g**), "AAI (high 3x)" (9 M) (**h**), "AAI (high 3x)" (12 M) (**i**), "AAI (high 14x)" (9 M and 12 M) (**j**), "AAI (3x) + $CCl_4$ " (9 M) (**k**), "AAI (3x) + $CCl_4$ " (12 M) (**l**), AAI (14x) + $CCl_4$ (6 M) (**m**), AAI (14x) + $CCl_4$ (9 M) (**n**), "AAI (7x)" (**o**), and "AAI (high 14x, *Pten*$^{LKO}$)" (**p**). (**q**) Kaplan–Meier plots of overall survival for these groups showing the occurrence of death and sacrifice at the age of 9 and 12 months. The observation time was from the end of administration to sacrifice. (**r**) Representative images (scale bars, 100 μm) of PicroSirius Red histochemistry of the liver in the different administration and control groups. (**s**) Gross appearance of hydronephrosis and renal cysts (scale bars, 1 cm) in the AAI alone administration groups. The fractions stand for the number of abnormal kidneys/number of total kidneys in each group. Asterisks signify significant differences using the two-sided Student's *t*-test or Wilcoxon rank-sum test and Fisher's exact test. $^{*}P < 0.05$; $^{**}P < 0.01$; NS, not significant.

**Supplementary Figure 2. AA can cause liver DNA damage.**

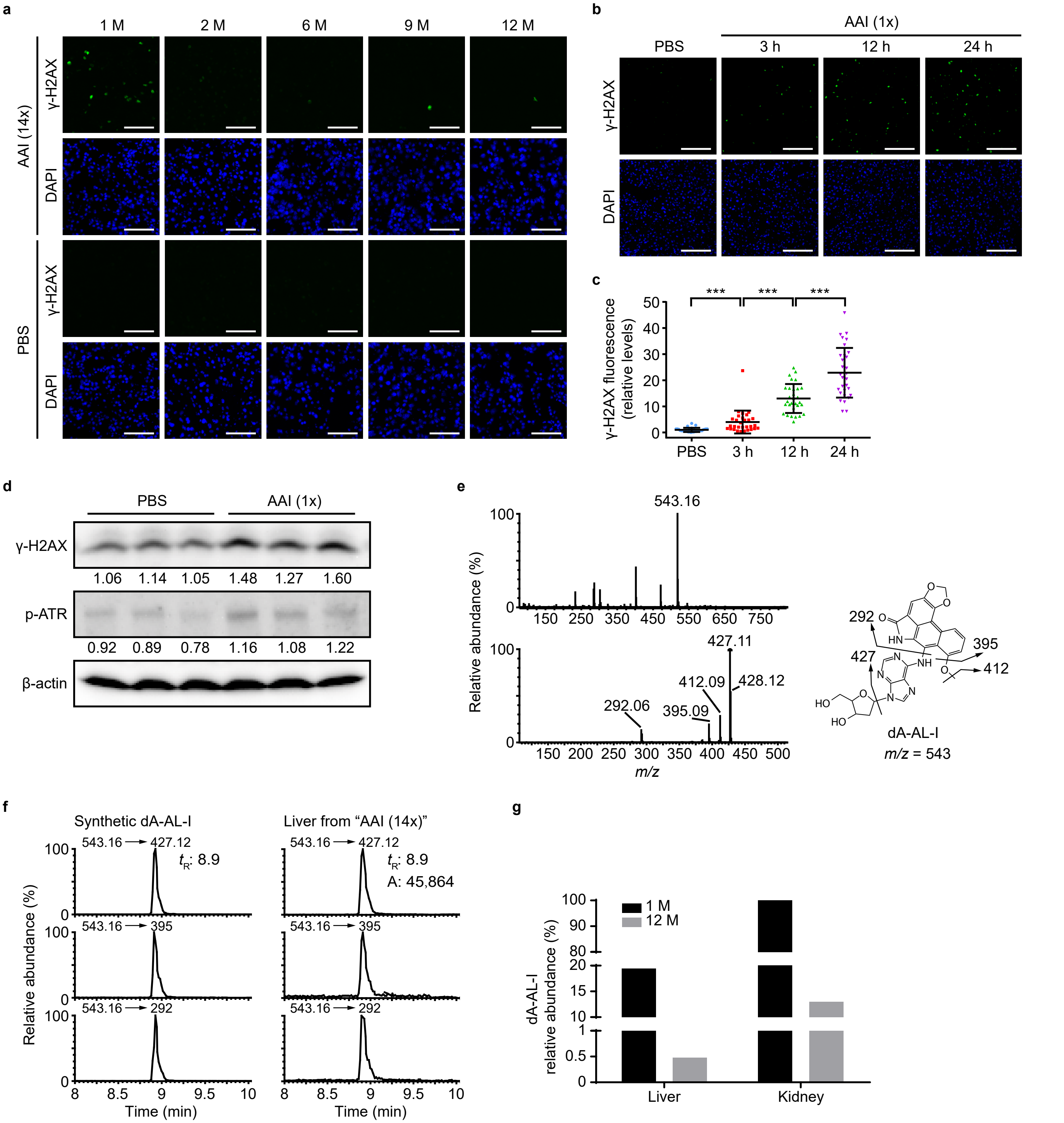


(**a**) γ-H2AX levels were measured by immunofluorescence assay in liver from “AAI (14x)” mice and the control group (PBS) at the indicated ages. (**b**, **c**) γ-H2AX levels were measured by immunofluorescence assay in liver from 2-week-old mice ($n$ = 3) at the indicated times after PBS or AAI (2.5 mg/kg) injection once, including representative images (**b**), and quantitative analyses (**c**). Values indicated by long and short horizontal lines represent the mean ± SD. (**d**) γ-H2AX and p-ATR (Ser428) levels were measured by the Western blotting assay ($n$ = 3 mice) in livers from 2-week-old mice at 12 h after PBS or AAI (2.5 mg/kg) injection once. (**e**) Ion spectra of synthetic dA-AL-I (*m/z* 543) acquired at the MS (up) and MS/MS (down) scan stage. (**f**) Extracted ion chromatograms of synthetic dA-AL-I and dA-AL-I in mouse samples. The relative abundance of dA-AL-I in livers and kidneys in the “AAI (14x)” model was measured at 543.16 → 427.12. (**g**) The relative abundance (*m/z* 427) of dA-AL-I was measured by MS in livers and kidneys in “AAI (14x)” mice at the indicated ages ($n$ = 1). Scale bars, 100 μm. Asterisks signify significant differences using the two-sided Student’s *t*-test or Wilcoxon rank-sum test. ***$P < 0.001$.

**Supplementary Figure 3. Mutational signatures of AAI-induced mouse liver cancer.**

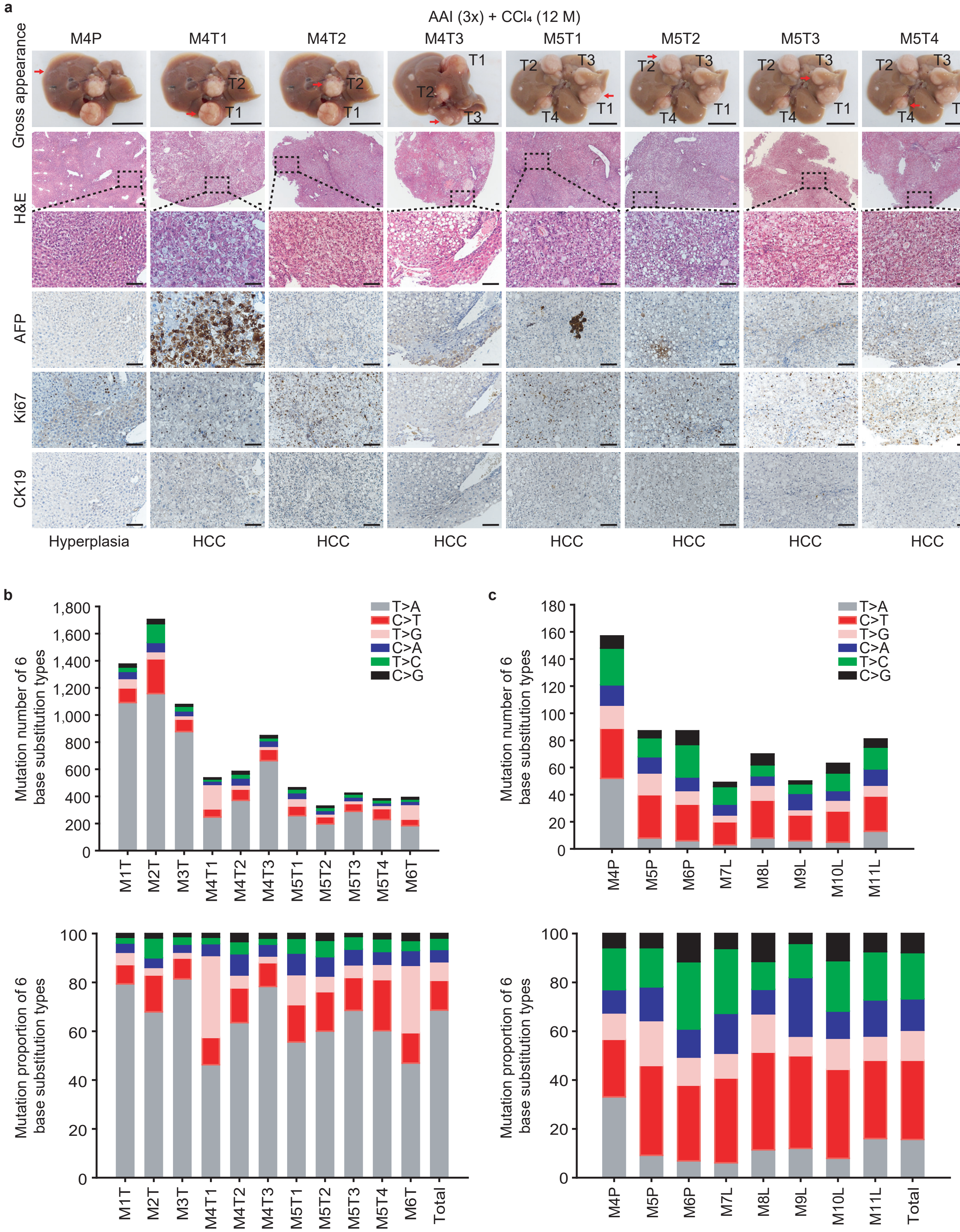

d

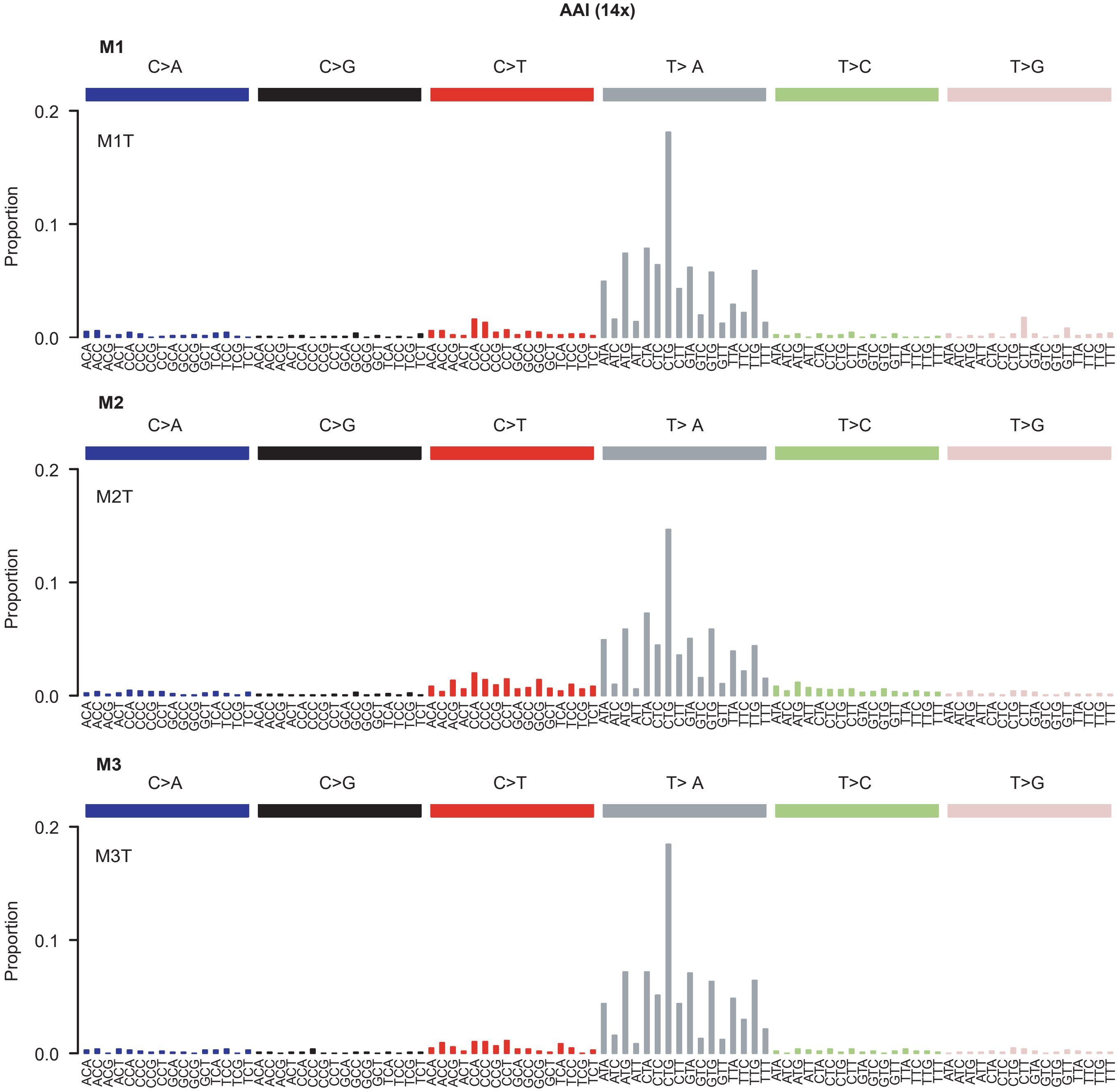
AAI (14x)
M1
C>A
C>G
C>T
T> A
T>C
T>G
M1T
Proportion
0.2
0.1
0.0
M2
M2T
M3
M3T

e

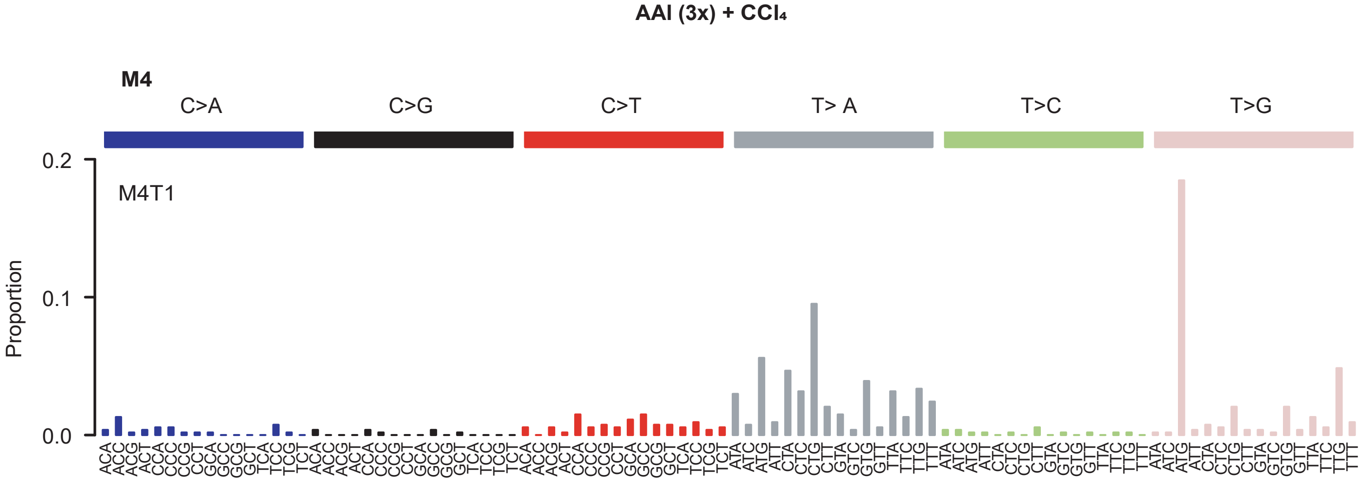
AAI (3x) + CCl₄
M4
C>A
C>G
C>T
T> A
T>C
T>G
M4T1
Proportion
0.2
0.1
0.0

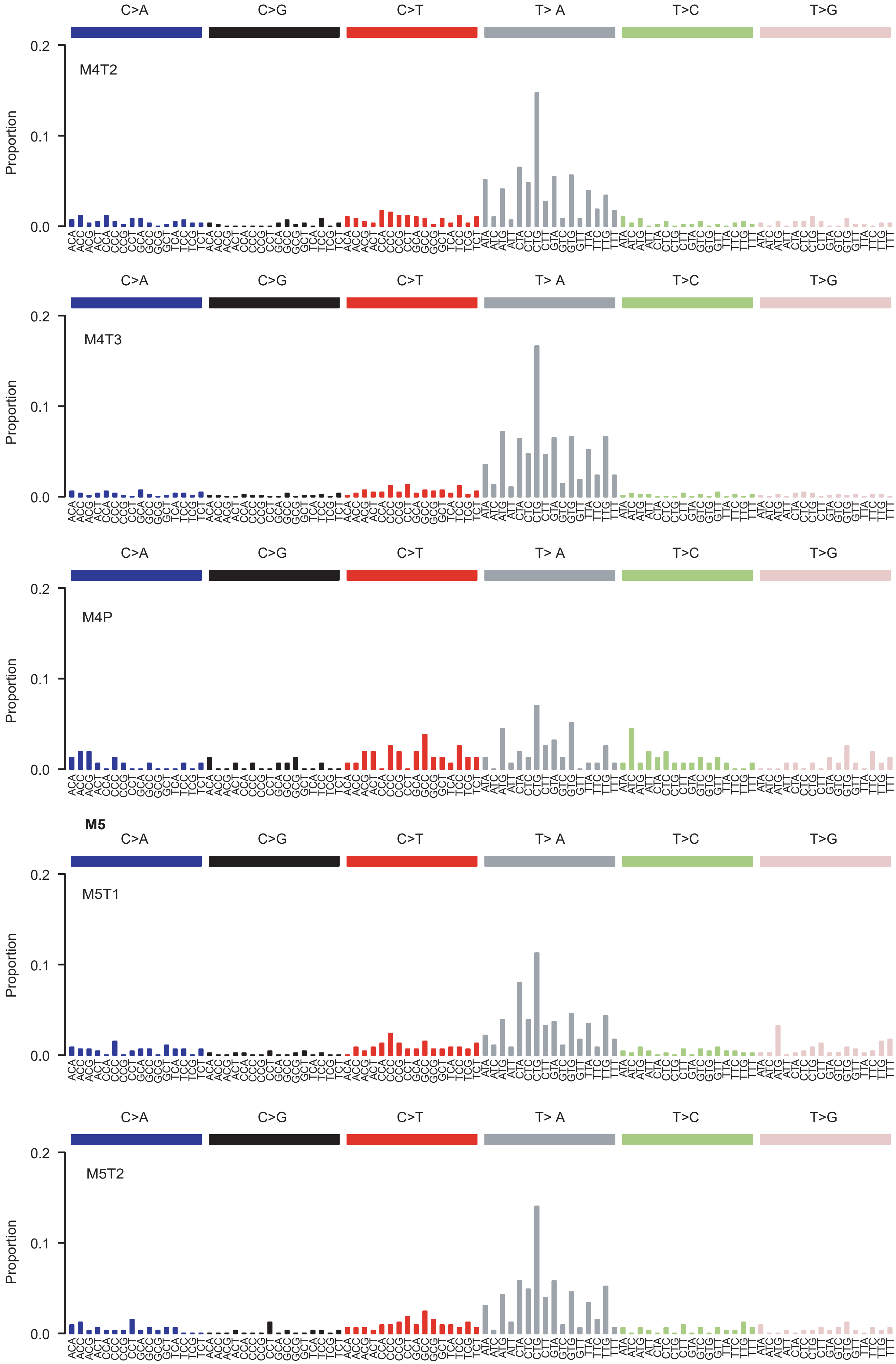
C>A
C>G
C>T
T> A
T>C
T>G
0.2
0.1
0.0
Proportion
M4T2
ACA ACC ACG ACT CCA CCC CCG CCT GCA GCC GCG GCT TCA TCC TCG TCT
ATA ATC ATG ATT CTA CTC CTG CTT GTA GTC GTG GTT TTA TTC TTG TTT
M4T3
M4P
M5
M5T1
M5T2

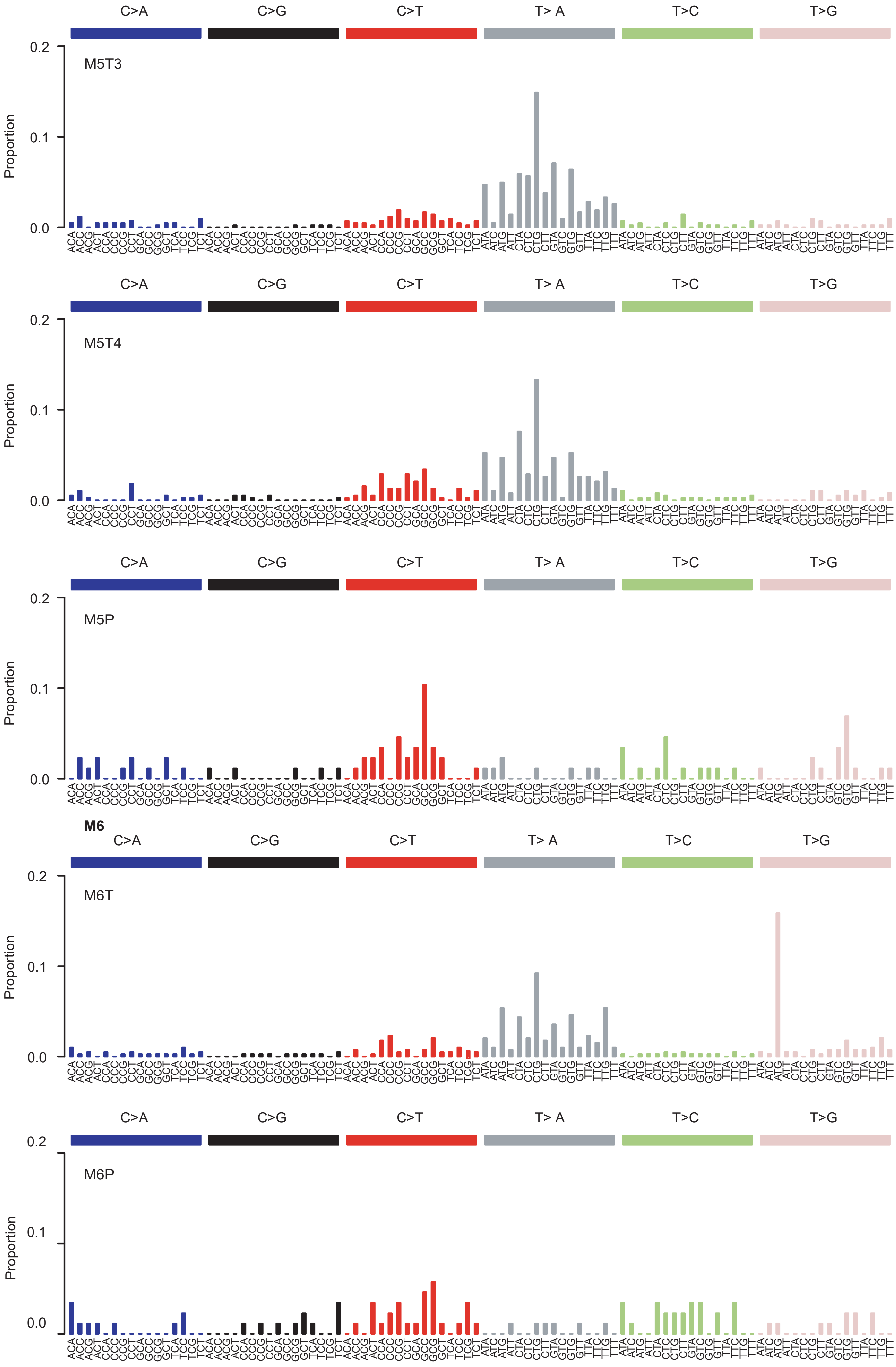
C>A
C>G
C>T
T> A
T>C
T>G
M5T3
M5T4
M5P
M6
M6T
M6P
Proportion
0.2
0.1
0.0
ACA ACC ACG ACT CCA CCC CCG CCT GCA GCC GCG GCT TCA TCC TCG TCT
ATA ATC ATG ATT CTA CTC CTG CTT GTA GTC GTG GTT TTA TTC TTG TTT

f

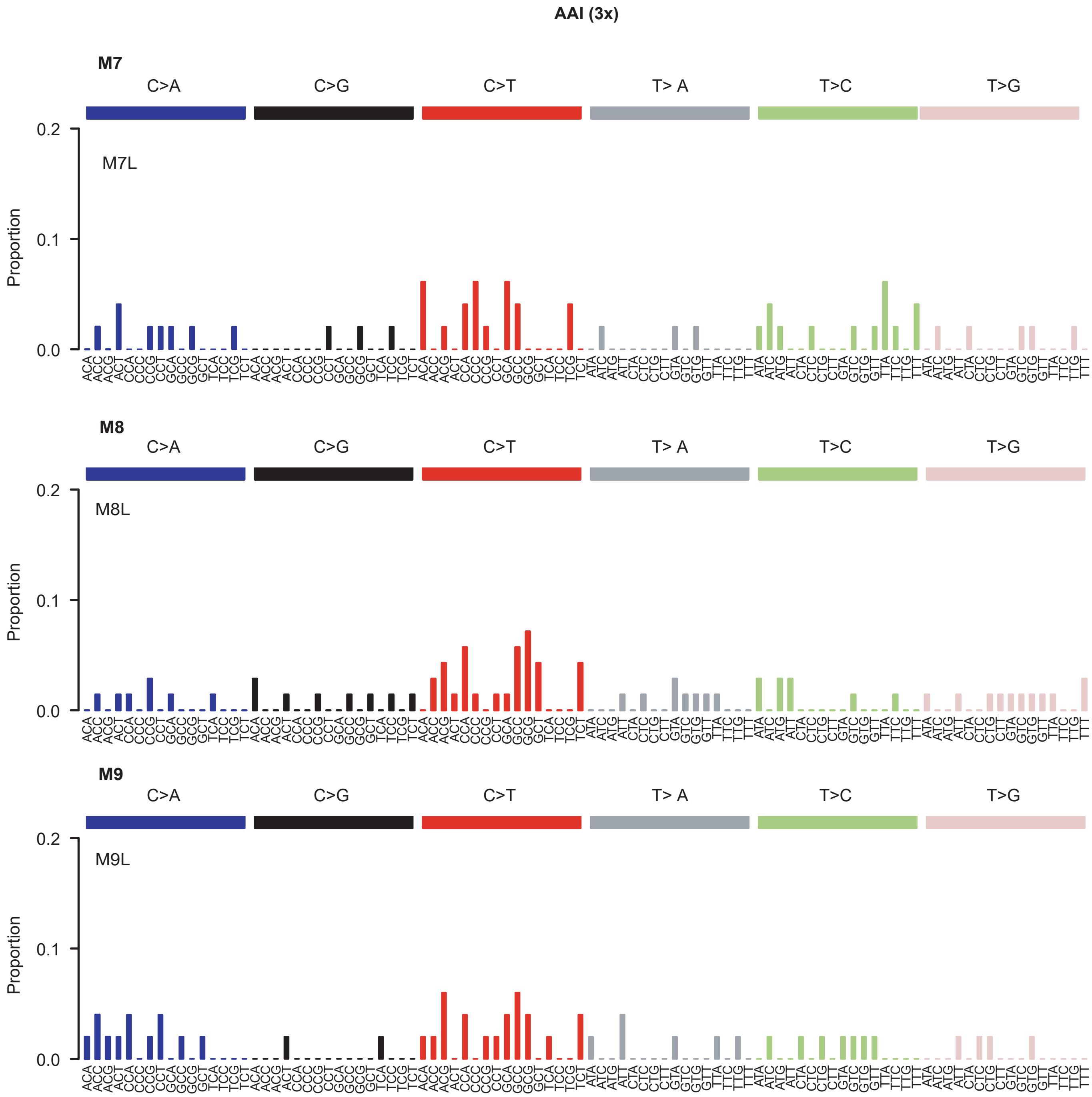

AAI (3x)
M7
C>A
C>G
C>T
T> A
T>C
T>G
M7L
Proportion
0.2
0.1
0.0
M8
C>A
C>G
C>T
T> A
T>C
T>G
M8L
Proportion
0.2
0.1
0.0
M9
C>A
C>G
C>T
T> A
T>C
T>G
M9L
Proportion
0.2
0.1
0.0


g

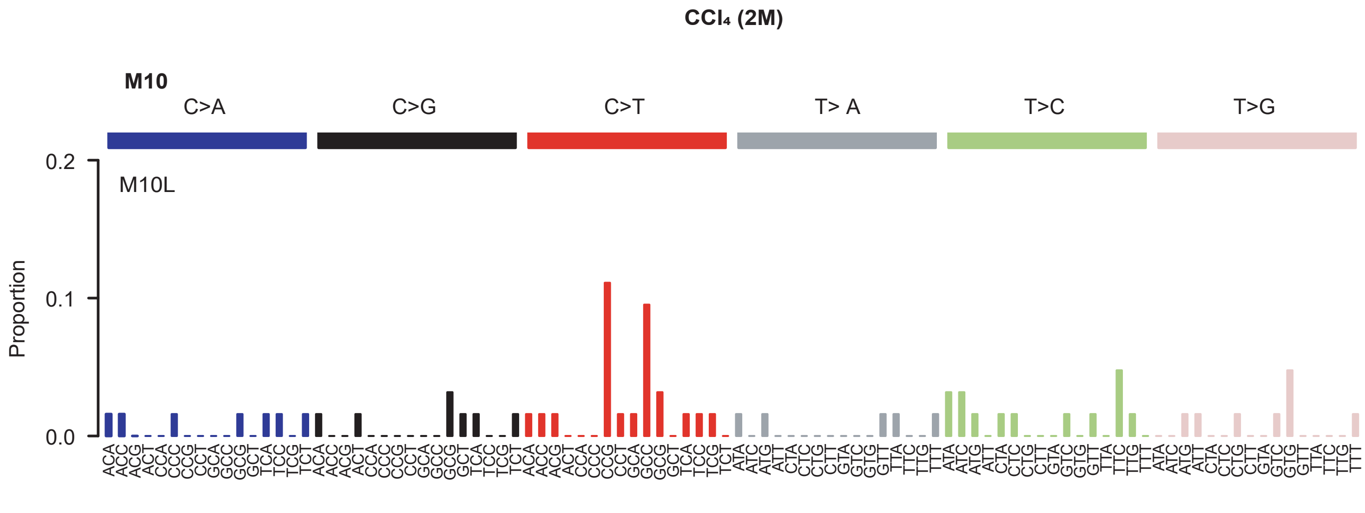

CCl₄ (2M)
M10
C>A
C>G
C>T
T> A
T>C
T>G
M10L
Proportion
0.2
0.1
0.0

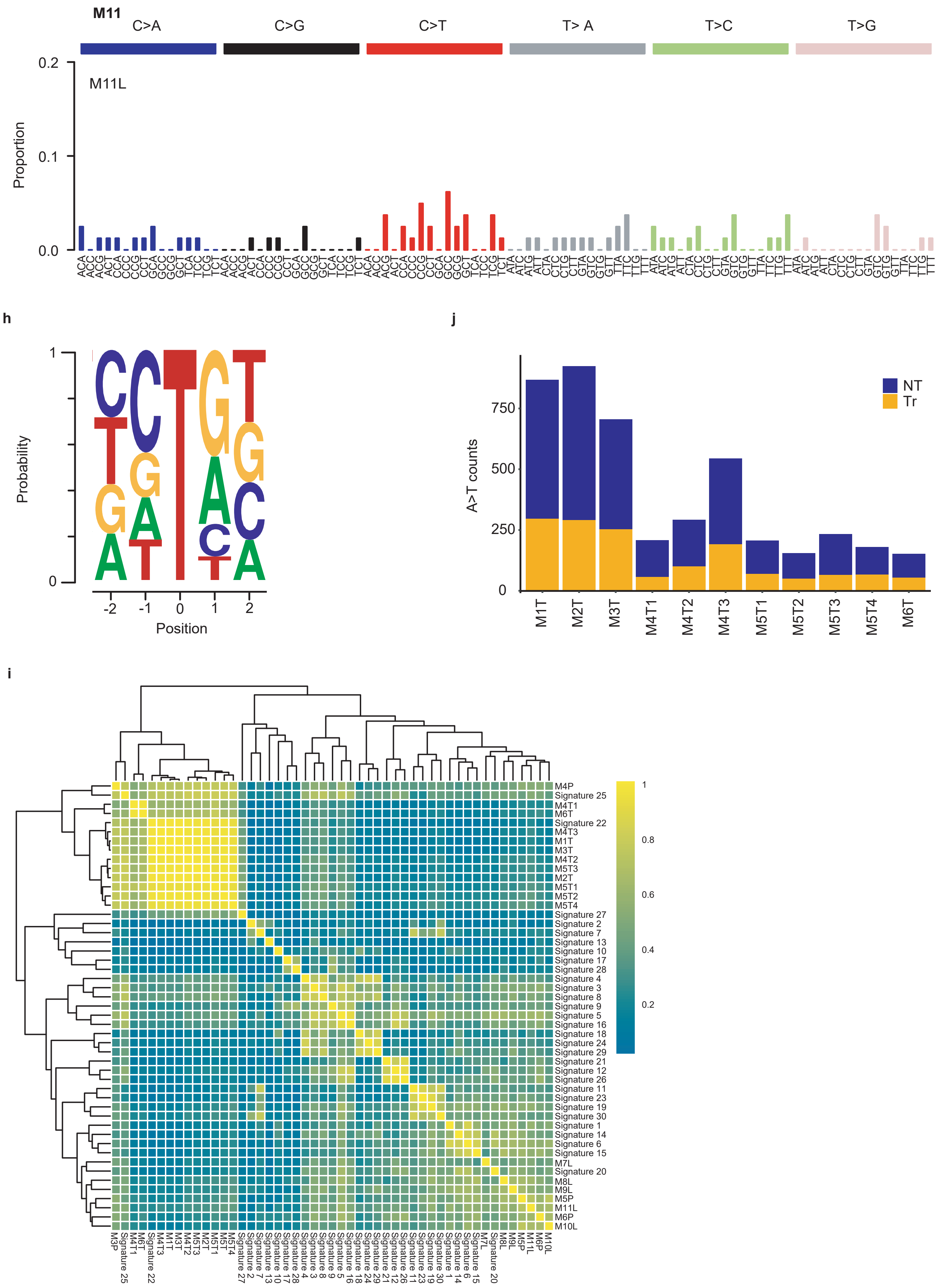
M11
C>A
C>G
C>T
T> A
T>C
T>G
M11L
Proportion
0.2
0.1
0.0
h
Probability
Position
j
A>T counts
NT
Tr
M1T
M2T
M3T
M4T1
M4T2
M4T3
M5T1
M5T2
M5T3
M5T4
M6T
i

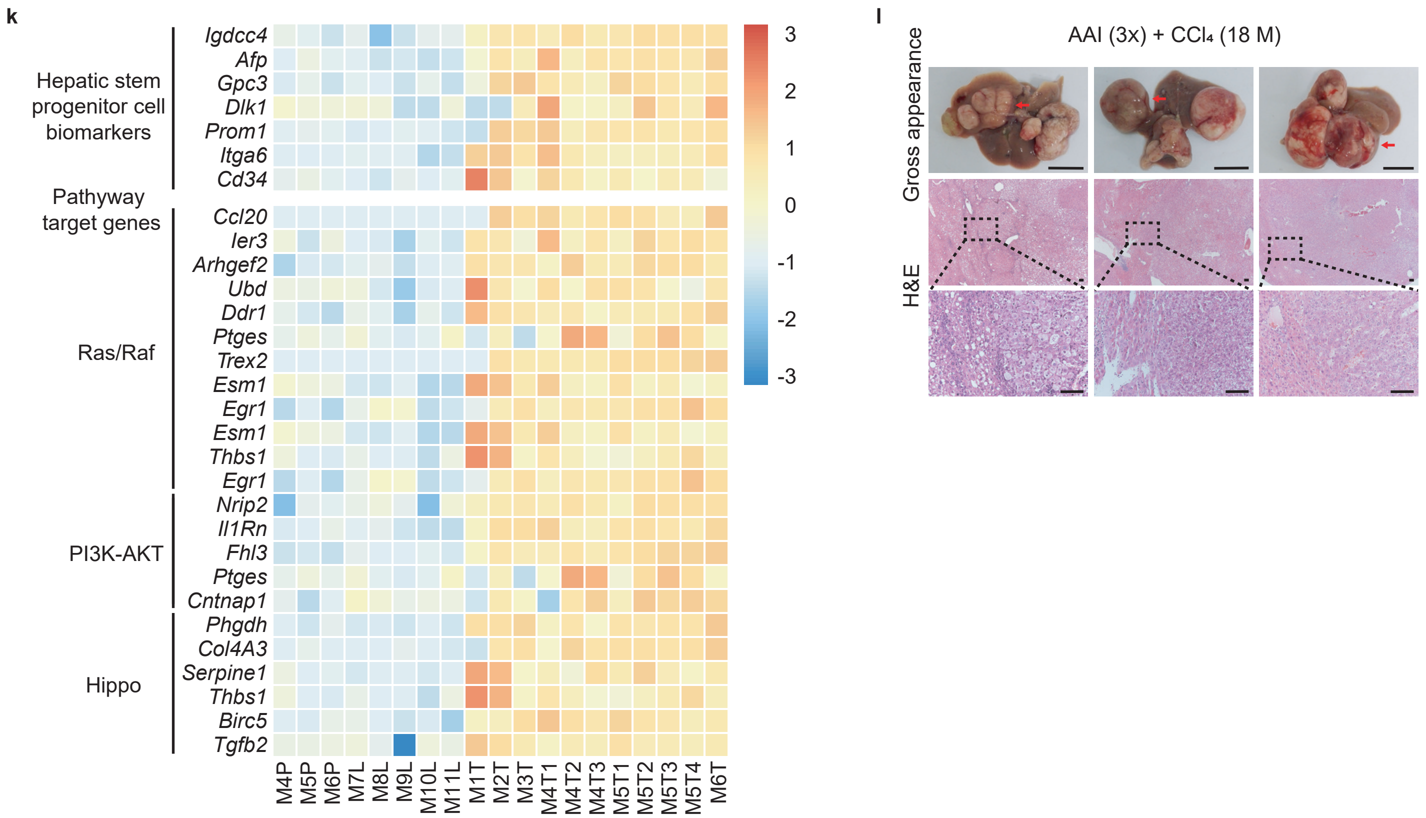


(**a**) Gross appearance (scale bars, 1 cm), H&E staining (scale bars, 100 μm), and IHC analysis with anti-AFP, Ki67 and CK19 antibodies (scale bars, 100 μm) of multiple tumor nodules in two mice (M4 and M5) from the "AAI (3x) + $CCl_4$" group. These samples were used for DNA sequencing. (**b**, **c**) Number and proportion of 6 base substitution types of AA-induced HCCs. (**d**-**g**) Mutational spectra of AA-induced HCCs and nontumor liver tissues. (**h**) Pentanucleotide sequence motif of the T>A mutations in mouse liver tumors. (**i**) Cosine similarities between the mutational spectra of AAI-induced HCCs, nontumor liver tissues and COSMIC mutational signatures. (**j**) Number of A>T mutations measured on the nontranscribed and transcribed strand in each mouse liver tumor. NT, nontranscribed strand; Tr, transcribed strand. (**k**) Heatmap of the top 30 differentially expressed genes related to the liver stem cell markers and selected driver mutation-responsive signaling pathways. (**l**) Gross appearance (scale bars, 1 cm) and H&E staining (scale bars, 100 μm) from the "AAI (3x) + $CCl_4$" group. These samples were used for Western blotting assay in **Fig. 4e**.

**Supplementary Figure 4. Clonal architecture and phylogenetic reconstructions of AAI-induced mouse liver cancer.**

a

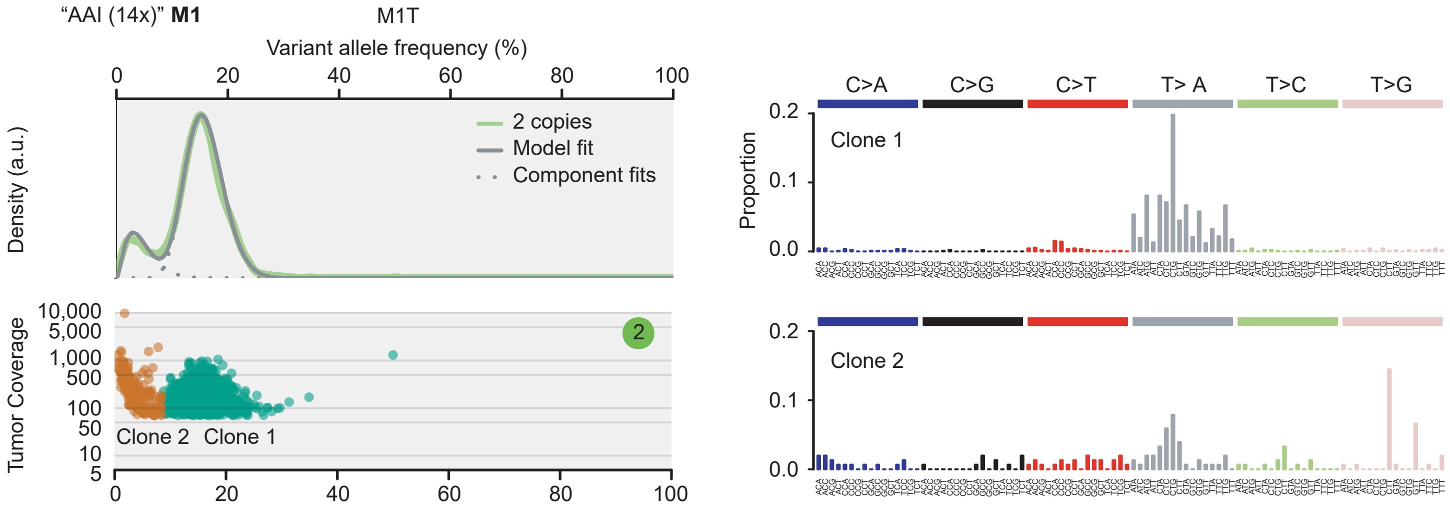


b

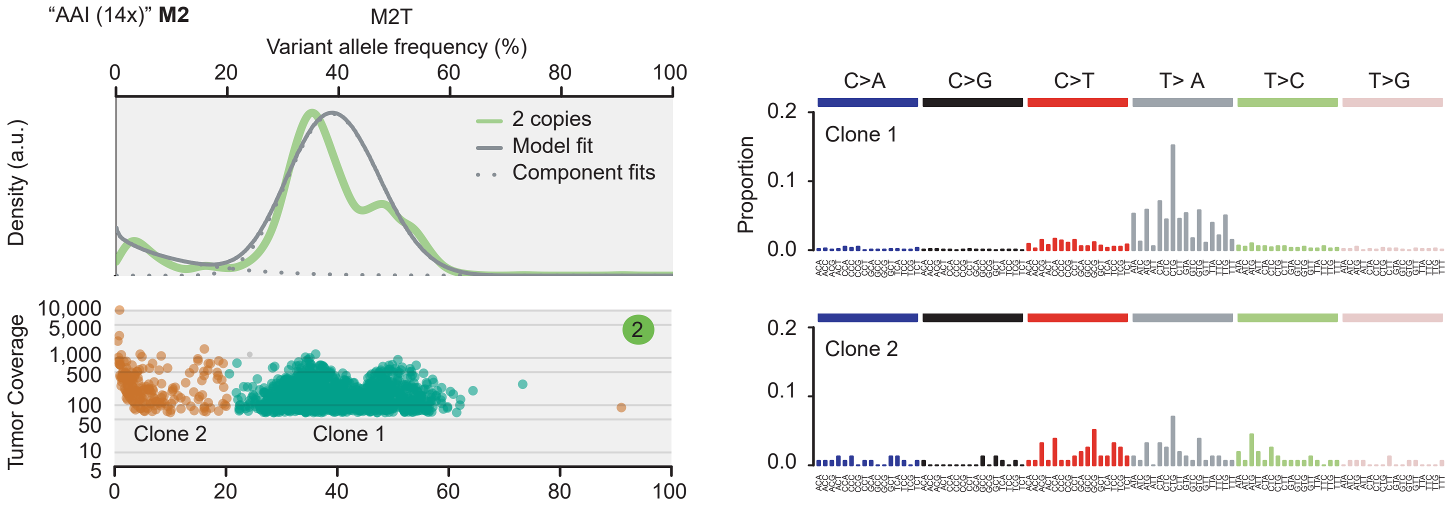


c

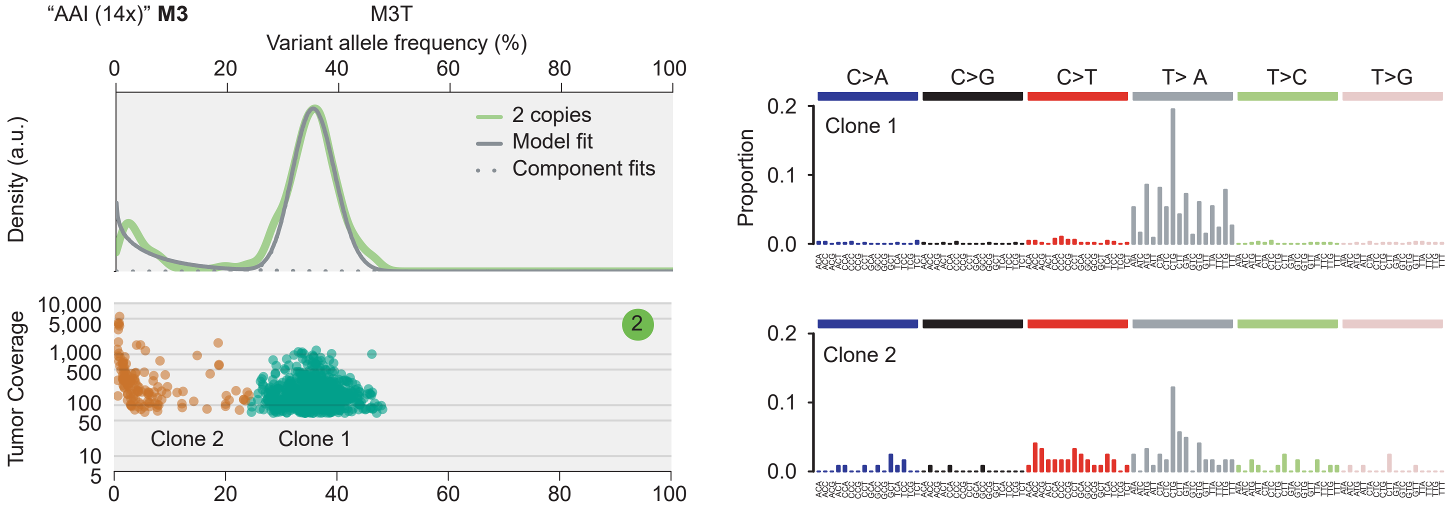

d

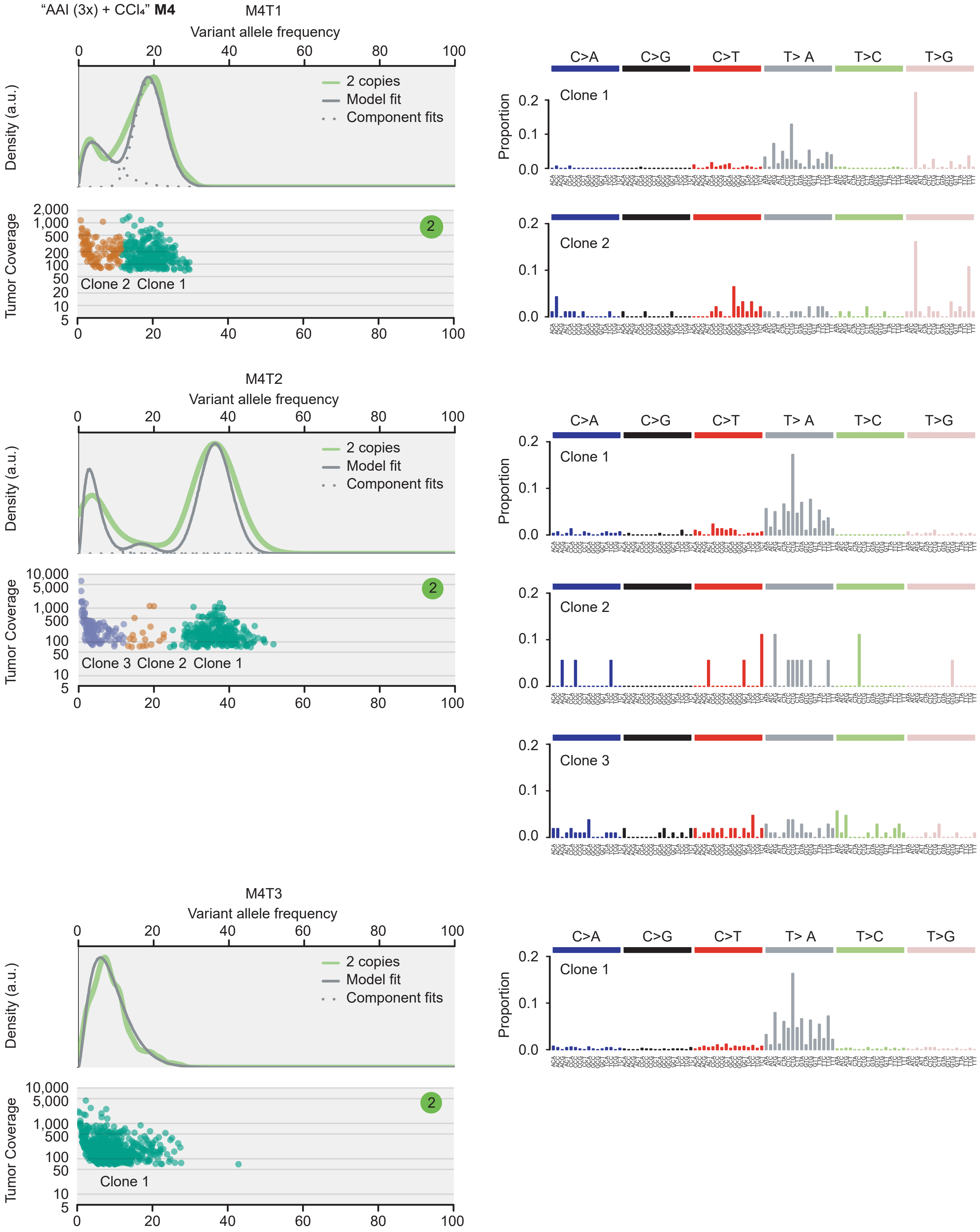

"AAI (3x) + CCl₄" M4
M4T1
Variant allele frequency
Density (a.u.)
2 copies
Model fit
Component fits
Tumor Coverage
Clone 2 Clone 1
C>A C>G C>T T> A T>C T>G
Proportion
Clone 1
Clone 2
M4T2
Variant allele frequency
Density (a.u.)
2 copies
Model fit
Component fits
Tumor Coverage
Clone 3 Clone 2 Clone 1
Clone 1
Clone 2
Clone 3
M4T3
Variant allele frequency
Density (a.u.)
2 copies
Model fit
Component fits
Tumor Coverage
Clone 1
Clone 1

e

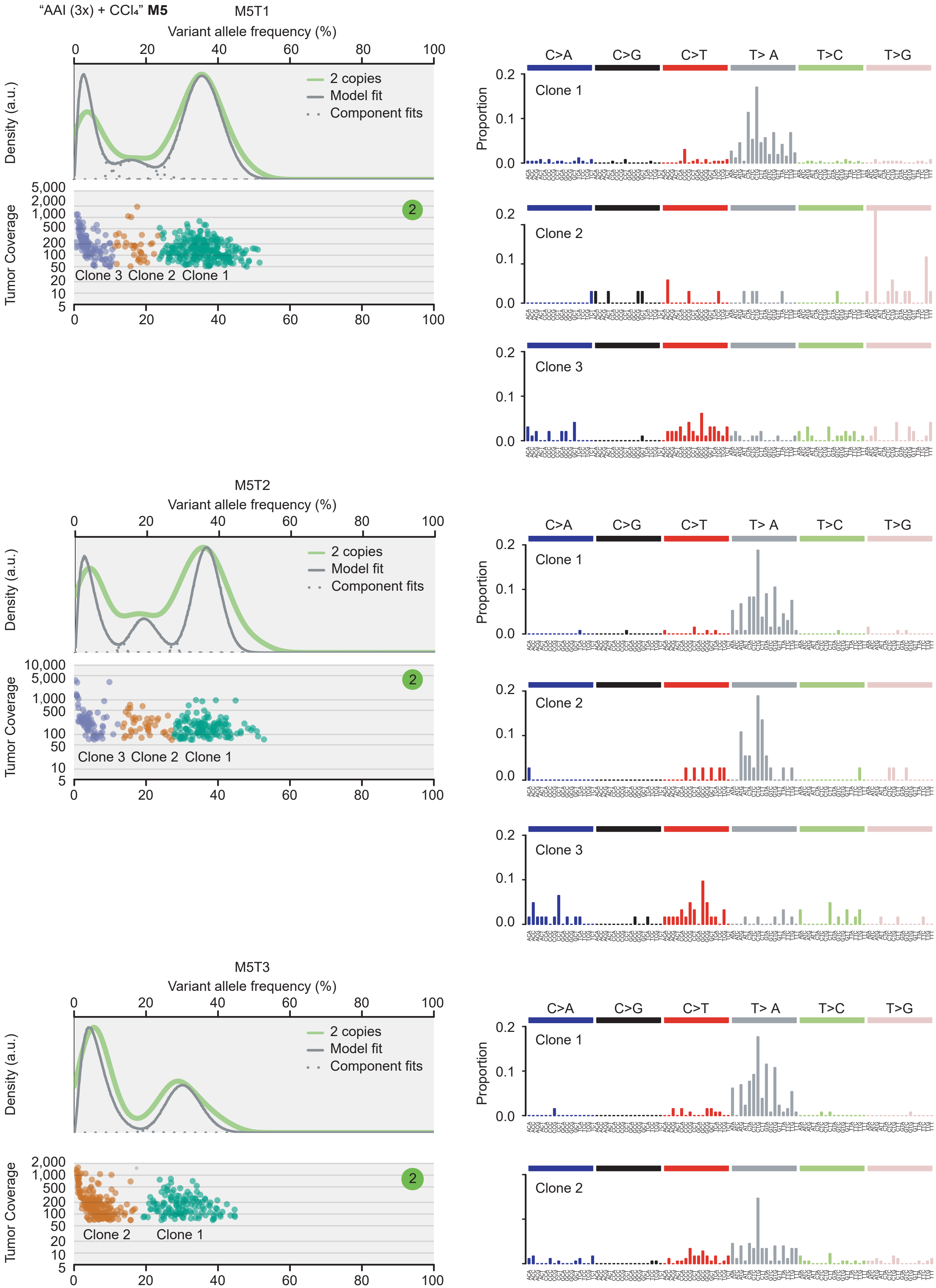
"AAI (3x) + $CCl_4$" **M5**
M5T1
Variant allele frequency (%)
0 20 40 60 80 100
Density (a.u.)
2 copies
Model fit
Component fits
Tumor Coverage
5,000 2,000 1,000 500 200 100 50 20 10 5
2
Clone 3 Clone 2 Clone 1
C>A C>G C>T T> A T>C T>G
Proportion
0.2 0.1 0.0
Clone 1
Clone 2
Clone 3
M5T2
Variant allele frequency (%)
0 20 40 60 80 100
Density (a.u.)
2 copies
Model fit
Component fits
Tumor Coverage
10,000 5,000 1,000 500 100 50 10 5
2
Clone 3 Clone 2 Clone 1
C>A C>G C>T T> A T>C T>G
Proportion
0.2 0.1 0.0
Clone 1
Clone 2
Clone 3
M5T3
Variant allele frequency (%)
0 20 40 60 80 100
Density (a.u.)
2 copies
Model fit
Component fits
Tumor Coverage
2,000 1,000 500 200 100 50 20 10 5
2
Clone 2 Clone 1
C>A C>G C>T T> A T>C T>G
Proportion
0.2 0.1 0.0
Clone 1
Clone 2

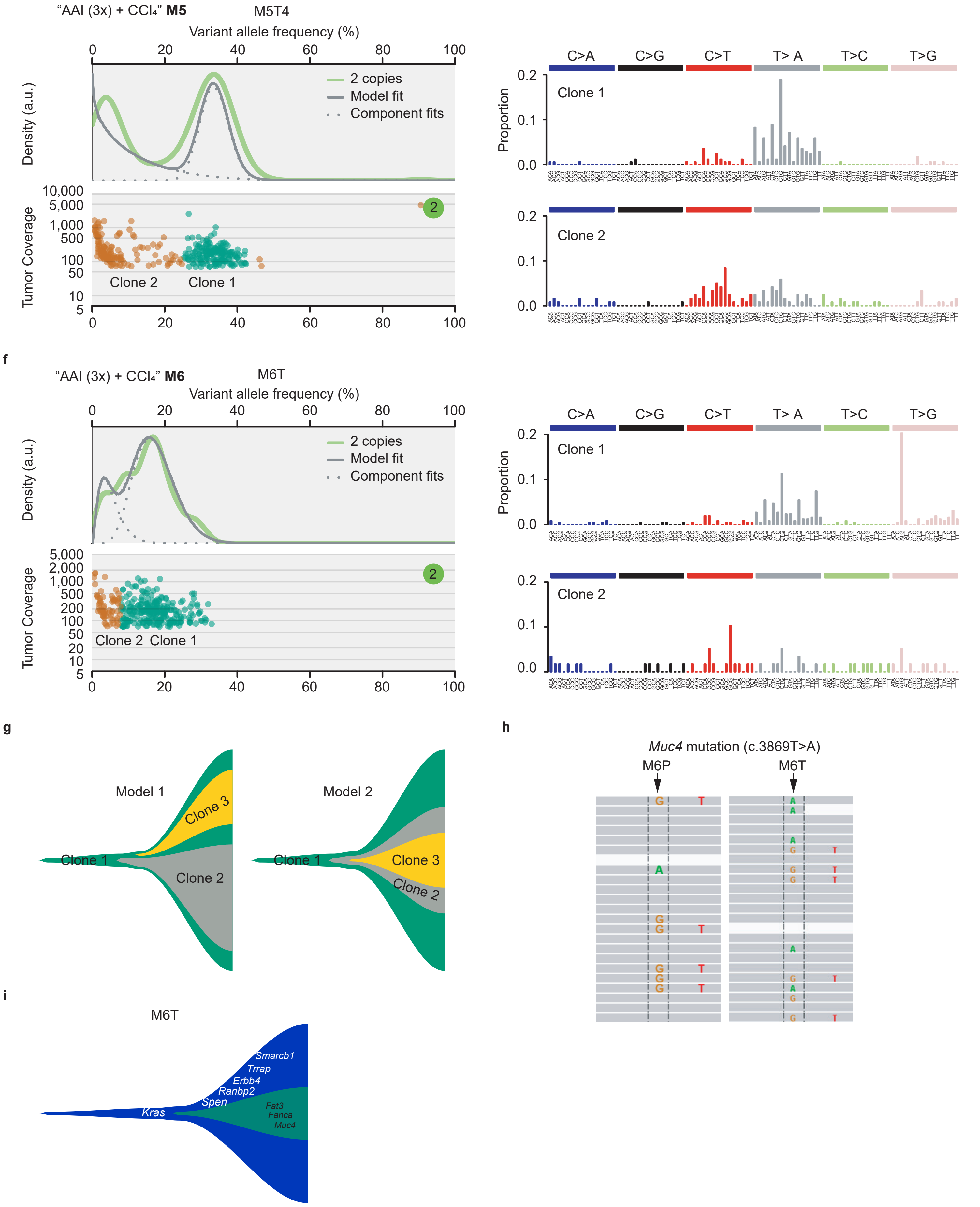


(**a**-**f**) Paired presentation of the clonal architecture (left) and the corresponding mutational profile (right) of AAI-induced mice liver tumors. (**g**) Tow models of the evolutionary trajectories of the tumors with three subclones. The left model assumes that the third weighted subclone forms independently from the second weighted subclone, whereas the right one assumes that the third weighted subclone arises within the second weighted clone. The left model is adopted by the illustration in **Fig. 5 e** and **f.** (**h**) M6T had more abundant *Muc4* mutated reads than in its paratumor tissue (M6P), as shown in the IGV of their mapped reads files. (**i**) Evolutionary model of M6T. The font size reflects the allele frequency of the genes in the tumor.

**Supplementary Figure 5. AA signatures in human liver cancer.**

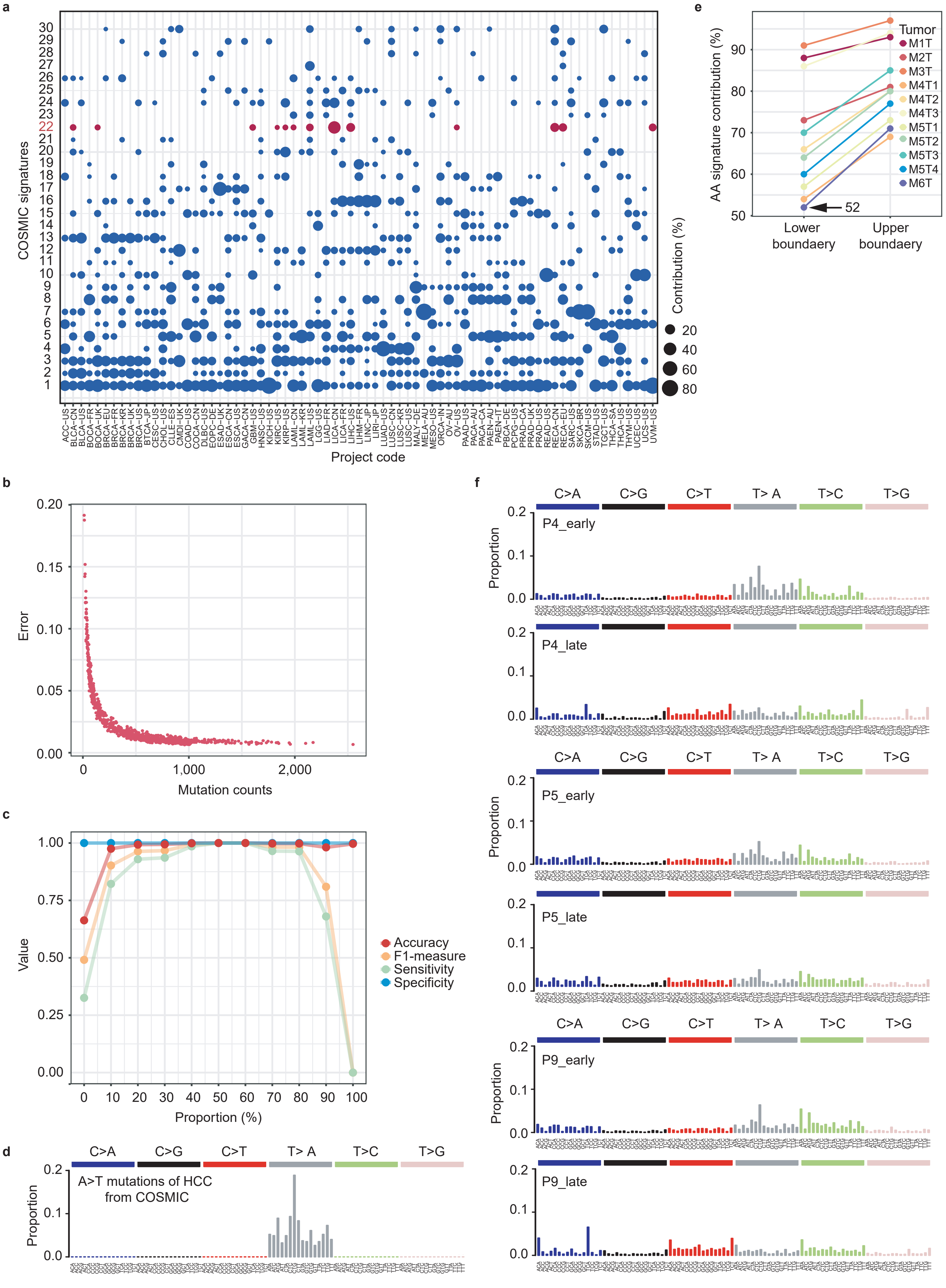

(**a**) COSMIC signature contributions in various TCGA and ICGC cancer projects using the webserver mSignatureDB. The AA signature (signature 22) was labeled in red. The sizes of the circles are in accordance with the contribution values. (**b**) The correlations between the mean square error (MSE) of the signature decomposing method and the mutation counts. (**c**) the accuracy, specificity, sensitivity and F1-measure evaluations for the bootstrapped signature decomposing method. (**d**) A>T transversion profile of liver cancer mutations in COSMIC. (**e**) The 95% confidential interval of the bootstrapped AA signature contributions in 11 mice liver tumors. M6T appeared to have the lowest lower boundary at 52%, which was adopted as one of the cutoffs for AA exposure evaluation in human liver cancers. (**f**) The mutational spectra of early and late assorted mutational profiles in Singapore AA-affected HCCs.
